# Neuron-specific protein network mapping of autism risk genes identifies shared biological mechanisms and disease relevant pathologies

**DOI:** 10.1101/2022.01.17.476220

**Authors:** Nadeem Murtaza, Annie A. Cheng, Chad O. Brown, Durga Praveen Meka, Shuai Hong, Jarryll A. Uy, Joelle El-Hajjar, Neta Pipko, Brianna K. Unda, Birgit Schwanke, Sansi Xing, Bhooma Thiruvahindrapuram, Worrawat Engchuan, Brett Trost, Eric Deneault, Froylan Calderon de Anda, Bradley W. Doble, James Ellis, Evdokia Anagnostou, Gary D. Bader, Stephen W. Scherer, Yu Lu, Karun K. Singh

## Abstract

There are hundreds of risk genes associated with autism spectrum disorder (ASD), but signaling networks at the protein level remain unexplored. We use neuron-specific proximity-labeling proteomics (BioID) to identify protein-protein interaction (PPI) networks for 41 ASD-risk genes. Neuron-specific PPI networks, including synaptic transmission proteins, are disrupted by *de novo* missense variants. The PPI network map reveals convergent pathways, including mitochondrial/metabolic processes, Wnt signaling, and MAPK signaling. CRISPR knockout reveal an association between mitochondrial activity and ASD-risk genes. The PPI network shows an enrichment of 112 additional ASD-risk genes and differentially expressed genes from post-mortem ASD patients. Clustering of risk genes based on PPI networks identifies gene groups corresponding to clinical behavior score severity. Our data reveal that cell type-specific PPI networks can identify individual and convergent ASD signaling networks, provide a method to assess patient variants, and reveal biological insight into disease mechanisms and sub-cohorts of ASD.

## Introduction

The risk of developing ASD has a strong genetic basis, including common and rare genetic variants (Gaugler et al., 2014; Iossifov et al., 2014; Robinson et al., 2016; Weiner et al., 2017). Large scale sequencing studies have identified hundreds of genes associated with ASD risk (Feliciano et al., 2019; Glessner et al., 2009; Ruzzo et al., 2019; Samocha et al., 2014; Satterstrom et al., 2020; Wilfert et al., 2021; Yuen et al., 2017). One hypothesis is multiple ASD risk genes converge in developmental brain signaling networks.

The majority of known convergent ASD-associated pathways are based on sequencing, transcriptomics, and gene co-expression analyses (Cederquist et al., 2020; Jin et al., 2020; Parikshak et al., 2013; Ramaswami et al., 2020; Sanders et al., 2015; Satterstrom et al., 2020; Willsey et al., 2013, 2021; Yuen et al., 2017). These studies have implicated synaptic transmission, translation, transcription, chromatin remodeling and gene splicing (Chang et al., 2015; O’Roak et al., 2012a; Ramaswami et al., 2020; Velmeshev et al., 2019; Voineagu et al., 2011). However, the majority of autism risk genes encode proteins, and protein-protein interactions (PPIs) are an essential mechanism of signaling (Neale et al., 2012; O’Roak et al., 2012b). Given that a large proportion of ASD genes have functions that do not occur in the nucleus (De Rubeis et al., 2014; Satterstrom et al., 2020), PPI networks provide an unbiased approach to gain insights into unknown convergent ASD disease processes (Kuzmanov and Emili, 2013; Murtaza et al., 2020). Previous studies have shown that ASD genes form PPI networks that can be shared (Chang et al., 2015; Chen et al., 2020; Corominas et al., 2014; Li et al., 2015; Neale et al., 2012; O’Roak et al., 2012a; Sakai et al., 2011). However, these data do not represent brain-specific networks (Lage, 2014). The lack of ASD risk-gene PPI networks represents a missing link towards understanding biological mechanisms in ASD.

Multiple proteomic techniques can be used to identify PPIs (reviewed in Richards et al., 2021). Further, many brain-expressed genes are large in size, including ASD-risk genes, which limits possible expression systems (Casanova et al., 2019). We balanced gene size limitations to capture strong and transient interactions, as well as proteins in close proximity, to build BioID PPI networks for ASD risk genes. We developed an *in vitro* proximity-labeling proteomics (BioID2) system with mouse primary neurons. Proximity-labeling proteomics can capture physiologically relevant interactomes in neural cell-types (Hamdan et al., 2020; Loh et al., 2016; Spence et al., 2019; Uezu et al., 2016) or to map cellular compartments (Go et al., 2021; Hung et al., 2017; Markmiller et al., 2018; Youn et al., 2018). We captured PPI networks from mouse cortical neurons given their role in ASD pathology, while growing them with glial cells to promote maturation (Barres, 2008; Stogsdill and Eroglu, 2017; Wilton et al., 2019).

In the current study, we screened the interactome of 41 ASD-risk proteins in neurons using BIOID. We targeted non-nuclear proteins (e.g., cytosolic proteins, receptors, kinases, and intracellular signaling proteins) and identified 1770 protein-level connections and neighbourhood proteins, which was approximately 50-times that reported in the STRING database (Snel et al., 2000). Convergent protein networks included synaptic transmission, mitochondrial/metabolic processes and Wnt signaling. Further investigation revealed that a subset of ASD genes regulates mitochondrial function. We also determined that *de novo* missense variants in synaptic or poorly characterized ASD-risk genes disrupt PPIs that lead to synaptic deficits. The shared ASD-risk gene network revealed an enrichment of an additional 112 ASD risk genes, and is enriched with ASD-associated brain cell types (Feliciano et al., 2019; Ruzzo et al., 2019; Sanders et al., 2015; Satterstrom et al., 2020; Yuen et al., 2017). We also found that individuals with variants within the 41-risk genes, with a high degree of shared interactions, had similar adaptive behavior scores using human clinical data from the MSSNG database (Trost et al., 2020; Yuen et al., 2017).

Taken together, we demonstrate that neuron-specific PPI networks provide a scalable approach to reveal individual and convergent disease mechanisms in ASD. This PPI network resource and screening system can be applied more broadly to additional autism risk genes to identify disease mechanisms that are not captured with current approaches.

## Results

### Development of a neuronal proximity-based proteomic system to identify PPI networks

We used mouse cortical neurons and glia co-cultures infected with neuron-specific human Synapsin1 promoter-driven lentiviral constructs expressing BioID2 fusion proteins (pLV-hSyn-tGFP-P2A-POI-13xLinker-BioID2-3xFLAG) (Figure 1A and Figure S1A). A 13x Gly-Ser linker sequence attached to the proteins-of-interest (POIs) increased the range of biotinylation to include direct/indirect interactors and proteins in close proximity. Embryonic age 16-17 (E16-17) mouse cortical neurons were infected at days *in vitro* (DIV) 14 (Figure 1A). Biotin was added at DIV17 for 18 hours, followed by processing for mass spectrometry. We used a Luciferase-P2A-BioID2-3xFLAG construct as a negative control (Figure S1B). To promote high efficiency infections, we optimized lentiviral production for small and large risk genes (Figure 1B).

**Figure 1.**
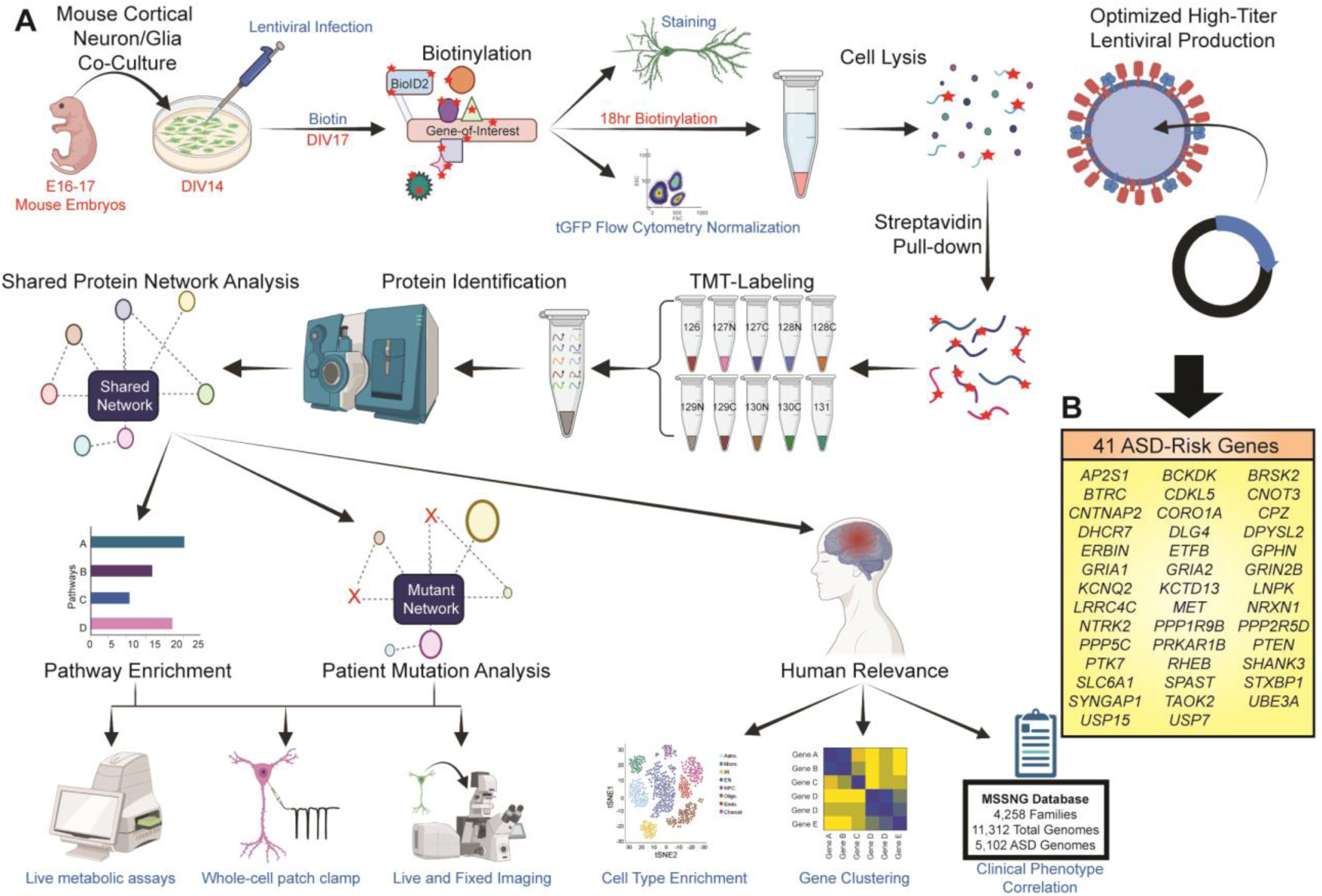
Development of a proximity-based proteomics screen to identify neuronal PPI networks for ASD-risk genes. (A) Workflow of neuron-specific BioID2 screen for identification of ASD-risk gene PPI networks. (B) List of 41 ASD-risk gene used in the BioID2 screen

To validate the BioID2 screening system, we used the well characterized excitatory synapse protein DLG4 (PSD95). Neurons expressing PSD95-BioID2 displayed punctate localization of BioID2-3xFLAG fusion proteins and biotinylated proteins around the dendrites, suggesting appropriate synaptic expression and biotinylation (Figure S1C). Synaptic punctate localization was also seen for TAOK2β-BioID2 as expected (Figure S1C) (Richter et al., 2019). The Luciferase-P2A-BioID2 control showed non-specific localization and biotinylation throughout the neuron as expected (Figure S1C). We identified 74 proteins that interact with PSD95, and Reactome pathway enrichment revealed neurotransmitter receptors and glutamatergic synapses (Figure S1D and Table S1). Comparison of our PSD95 PPI network with previously published data revealed 25 shared proteins (Fernández et al., 2009; Uezu et al., 2016) (Figure S1E), highlighting that our system captures relevant networks, acknowledging there are differences between methods and approaches.

Cortical neurons are a major cell type associated with ASD (Satterstrom et al., 2020); however, BioID approaches have typically been done primarily in cell lines (Go et al., 2021). To determine the importance of using neurons for the screen, we performed BioID2 in HEK293 cells (Figure S2 and Table S2). The PSD95 PPI network from HEK293 cells showed an absence of synaptic pathways (Figure S2A). Furthermore, BioID2 of six ASD risk proteins in HEK293 cells revealed a significant loss of protein interactions localized in neuron-specific compartments, and large differences in the PPI network compared to mouse neurons (Figure S2B-S2G, and Table S3).

To further validate the neuron-specific BioID2 screening system, we targeted compartment proteins (Fazal et al., 2019), including the microtubules (MAP2C), endoplasmic reticulum network (CANX), plasma membrane (PDGFR transmembrane domain), trans-Golgi apparatus (TGOLN), the presynaptic terminal (SNCA), and nucleus (MECP2). Compartment analysis of each PPI network revealed enrichment of the compartments expected for each bait protein (Figure S3, Figure S4D, Table S1 and Table S4). SNCA did not have a strong enrichment of presynaptic compartments; however, it did identify pathways involving axons, growth cones and the synapse (Figure S3E). BioID2 of MECP2, a nuclear protein, indicated localization to the nucleus (Figure S4A) and interaction with proteins enriched in nucleus-specific pathways (Figure S4B). The MECP2 PPI network was different in mouse neurons compared to HEK293 cells (Figure S4C and S4D), suggesting that mouse neurons have differing MECP2 interactions localized to the nucleus. Further, the PPI network of MECP2 did not include some of the known protein interactions in mouse neurons (e.g., ATRX, CREB1, SIN3A, NCor, and TET1), possibly due to the presence of highly biotinylated endogenous nuclear proteins or experimental artifacts. The enrichment of proteins specific to each compartment analyzed provides validation that the BioID2 screen in mouse cortical neurons can provide relevant PPI networks.

### Identification of a shared PPI network map and common pathways of 41 ASD-risk genes in mouse cortical neurons

To develop a shared PPI network map for ASD risk genes, we selected 41 ASD-risk genes that encode proteins with a range of molecular functions (Figure 1B). These genes were chosen from a combined list of ASD-risk genes from the SPARK, SFARI category 1, 2, and syndromic gene lists and previous sequencing studies (Feliciano et al., 2019; Ruzzo et al., 2019; Sanders et al., 2015; Satterstrom et al., 2020; Wilfert et al., 2021; Yuen et al., 2017). The final list was filtered for size limitations of the lentivirus, however we included some large genes (>4kb) such as SHANK3 and SYNGAP1 by optimizing lentivirus production. All genes chosen for the screen have a cytoplasmic function or localization (Satterstrom et al., 2020). For each gene, the human cDNA was cloned into a BioID2 lentiviral backbone and protein expression was confirmed (Figure S5A). We identified the individual PPI networks and enriched Reactome pathways, biological processes and cellular compartments for each gene in mouse cortical neurons (Table S1 and Table S5). Validation of BioID protein hits verified the interactions between Taok2 and Fbxo41, similar to another publication (Pennemann et al., 2021) (Figure S5B). We also verified interactions between Fmrp and Stxbp1, Taok2 and Syngap1, and the mitochondrial citrate synthase protein (Cs) and Syngap1 (Figure S5C-S5E). Furthermore, we found that Psd95 and Taok2β, and to a lesser degree, Shank2 and Taok2β, co-localized in mouse cortical neurons (Figure S5F).

The 41 ASD-risk gene PPI network consisted of 1109 proteins (41 ASD bait proteins and 1068 prey proteins) and 2349 connections. Every ASD bait protein shared at least 4 shared prey proteins with one other ASD bait protein (Figure 2A). Reciprocal identification was observed between DLG4 and CDKL5, GRIA1, GRIA2, or SYNGAP1 and between GRIA1 and GRIA2. BioID2 of CDKL5, DLG4, LRRC4C, SYNGAP1, and TAOK2 identified the most ASD bait proteins, suggesting high connectivity between a subset of ASD bait proteins. We identified 3 groups of highly connected ASD-risk genes from the individual PPI networks of the 41 ASD-risk genes (Group 1, 2 and 3), based on the correlation between individual PPI networks (Figure 2B). Groups 1 and 2 showed high connectivity between the ASD-risk genes within each group, whereas connectivity was lower in Group 3.

**Figure 2.**
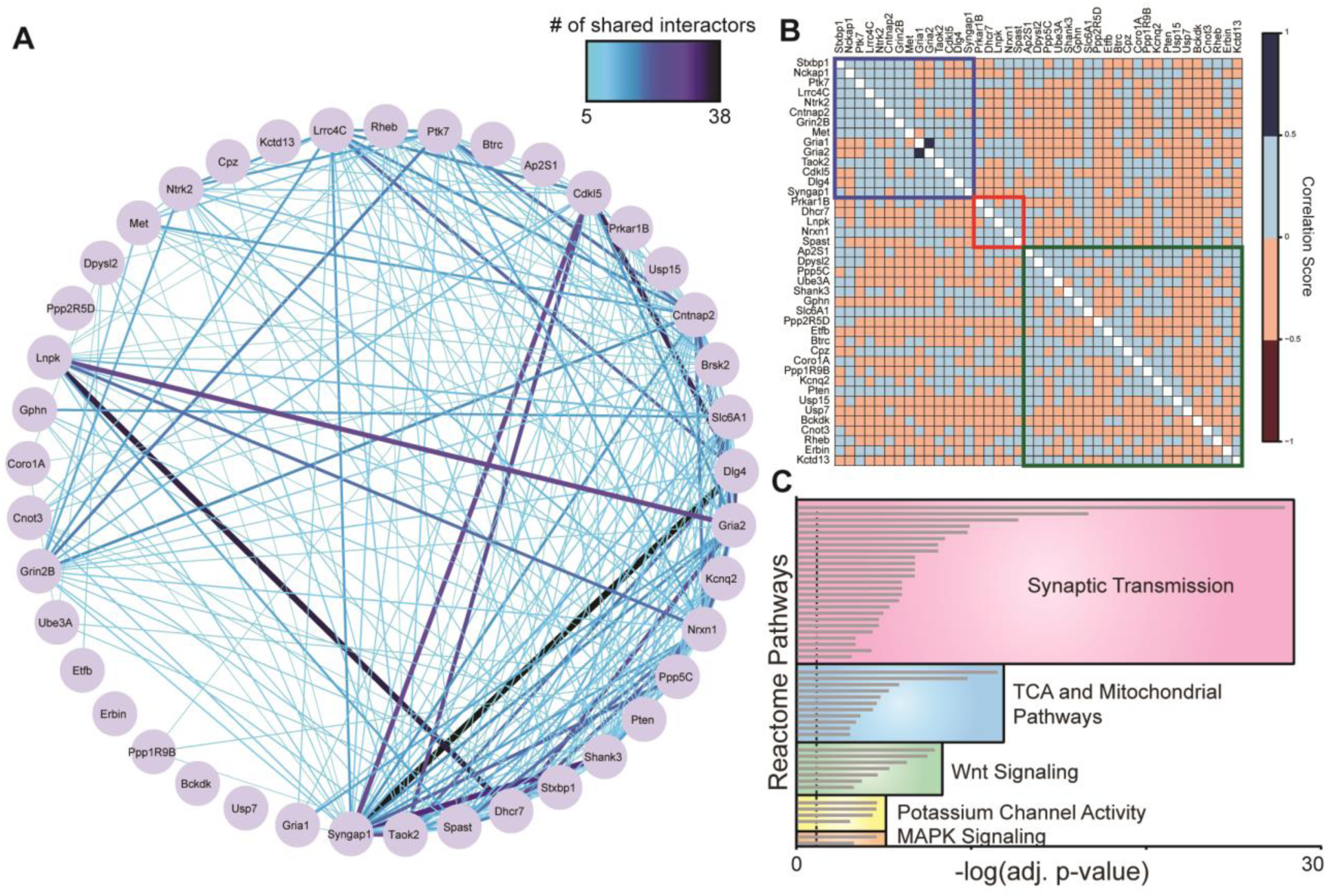
Neuron-specific PPI network map identifies convergent signaling pathways between 41 ASD-risk genes. (A) Shared PPI network map of 41 ASD-risk bait genes. Large red nodes represent bait proteins. Color and increased thickness of connecting lines represent number of interactions (direct or shared prey proteins) between bait genes. (B) Correlation plot of 41 ASD-risk genes through individual PPI networks. Genes were ordered by hierarchical clustering and clustered using kmeans (k = 3). (C) Top 50 enriched Reactome pathways of the shared 41 ASD-risk gene PPI network map. Individual pathways are grouped by functional similarity (g:Profiler, Bejamini-Hochberg FDR adj. p<0.05).

We compared our results with physical interactions between the 41 ASD bait proteins extracted from the STRING database (greater than or equal to medium confidence, 0.4). Our neuron-generated 41 ASD-risk gene PPI network had a near 50-fold increase in the number of connections compared to the STRING-db network (Figure 2A and S6A). STRING is primarily derived from non-neuronal sources using gene co-expression or direct interaction data (Lage, 2014), which do not capture proteins within close proximity to the POI. However, the PPI networks we identified include both direct interacting proteins, shared interacting proteins and close proximity proteins, highlighting potential connections missed by traditional methods.

The most significant pathways in the shared 41 ASD-risk gene PPI network involve synaptic transmission (Figure 2C and Table S6). Other enriched pathways included TCA cycle and mitochondrial activity, Wnt signaling, potassium channel activity, and MAPK signaling (Figure 2C and S6B, and Table S6). The majority of the shared ASD-risk PPI network localized to specific cellular compartments including axons, dendrites and synapses (Figure S6C and Table S6), while the majority of biological processes involve synaptic signaling and organization, and protein transport (Figure S6D and Table S6). Shared pathways in the ASD-risk gene PPI network reflect the major role of synaptic dysfunction in ASD, but also highlight that less well-studied mechanisms are important in ASD pathology.

### The shared PPI network map identifies the tricarboxylic acid (TCA) cycle and pyruvate metabolism as a common signaling pathway in ASD

The 41-gene shared PPI network enriched for the TCA cycle and pyruvate metabolism proteins, implicating dysregulation in mitochondrial function and cellular metabolism. This pathway has been associated with some ASD associated genes such as *Fmr1* and *Mecp2*, but it is unknown whether this extends to other ASD risk genes (Bülow et al., 2021; Licznerski et al., 2020; Shulyakova et al., 2017). Previous clinical studies have identified abnormal mitochondrial function in ASD patient lymphoblastoid cells (Frye, 2020; Rose et al., 2014, 2018; Shen et al., 2019a). The TCA cycle and respiratory electron transfer chain protein Reactome pathway was highly enriched in the shared ASD-risk gene PPI network map (adj. p-value = 3.14×10^-12^), even without the PPI network for the mitochondrial protein ETFB (adj. p-value = 3.21×10^-6^) (Table S6). 28 out of 41 ASD-risk genes associated with at least one TCA cycle and pyruvate metabolism associated protein (Figure 3A). Citrate synthase (CS), which is involved in turning acetyl-CoA into citrate early in the TCA cycle, interacted with eight ASD bait proteins (ERBIN, MET, NRXN1, SHANK3, SPAST, STXBP1, SYNGAP1, TAOK2). The TCA cycle and pyruvate metabolism are essential for proper cellular respiration (adj p-value = 1.64×10^-10^) (Table S6). Therefore, we investigated this finding using a gene not previously associated with mitochondrial and metabolic processes, *TAOK2*, a gene in the 16p11.2 deletion/duplication region associated with ASD (Calderon de Anda et al., 2012; Richter et al., 2019; Ultanir et al., 2014; Yadav et al., 2017). We measured cellular respiration using live-cell metabolic assays in mouse cortical neurons. *Taok2* heterozygous knockout (Het) mouse cortical neurons showed an increase in maximal respiration, proton leak, non-mitochondrial respiration, spare respiratory capacity, and a decrease in ATP coupling efficiency (Figure 3B and 3C, and Figure S7A-S7D) compared to wildtype (WT) neurons. These changes indicated less functional mitochondria, which was corroborated by proteomic analysis of post-synaptic density (PSD) fractions from *Taok2* WT and homozygous knockout (KO) mouse cortices (Figure S7E). *Taok2* KO mice PSD fractions had downregulation of proteins involved in synaptic function and activity, and ETC complex proteins (Figure S7F and Table S7). Analysis at the transcriptome level also revealed reduced mRNA levels of mitochondrial membrane proteins in *Taok2* KO mouse cortices (Figure S7G and S7H, and Table S7), coinciding with the reduced protein levels of mitochondrial proteins (Figure S7F). Further investigation revealed that *Taok2* Het and KO neurons have a reduction in active Teramethylrhodamine (TMRM)-stained mitochondria (Figure 3D and 3E, and Figure S7I). There was also an increase in mitochondria size and number using the marker TOMM20 (Figure 3F and 3G). We examined the morphology of mitochondria *in vivo* from electron microscopy (EM) images from WT and *Taok2* KO mouse brains (Richter et al., 2019). *Taok2* KO mouse neurons had altered mitochondrial morphology with a reduction in category 1 and 3 mitochondria (typical morphology), and an increase in category 2 mitochondria at synapses (Figure 3H and 3I) (Da Costa et al., 2018). Category 2 mitochondria indicate enlarged non-contiguous mitochondrial cristae, which can reduce oxidative phosphorylation and cause aberrant mitochondrial protein translation (John et al., 2005; Schmidt et al., 2010). We extended these studies to human induced pluripotent stem cell (iPSC)-derived NGN2-neurons. We generated isogenic *TAOK2* KO and heterozygous knock-in *TAOK2 A135P* iPSC lines. A135P is a *de novo* missense variant which renders TAOK2 as kinase dead (Richter et al., 2019). We generated human neurons through direct differentiation of iPSCs via NGN2 overexpression and found altered cellular respiration in *TAOK2* KO neurons, similar to mouse neurons, and increases in the spare respiratory capacity of *TAOK2* KO and *A135P* neurons (Figure S7J-S7L). *TAOK2* KO and *A135P* human neurons transfected with Mito7-DsRed also displayed an increase in mitochondrial puncta size, suggesting an increase in the number or size of the mitochondria (Figure S7M and S7N). To determine if these changes were due to long-term deficits caused by loss of TAOK2 function, we used acute *Taok2* shRNA knock-down and found a decrease in mitochondrial membrane potential (Figure S8A-S8C). Taken together, using *TAOK2* as a validation gene, we determined that mouse and human models with disruption of *TAOK2* have altered cellular respiration, likely caused by altered activity, size and number of mitochondria.

**Figure 3.**
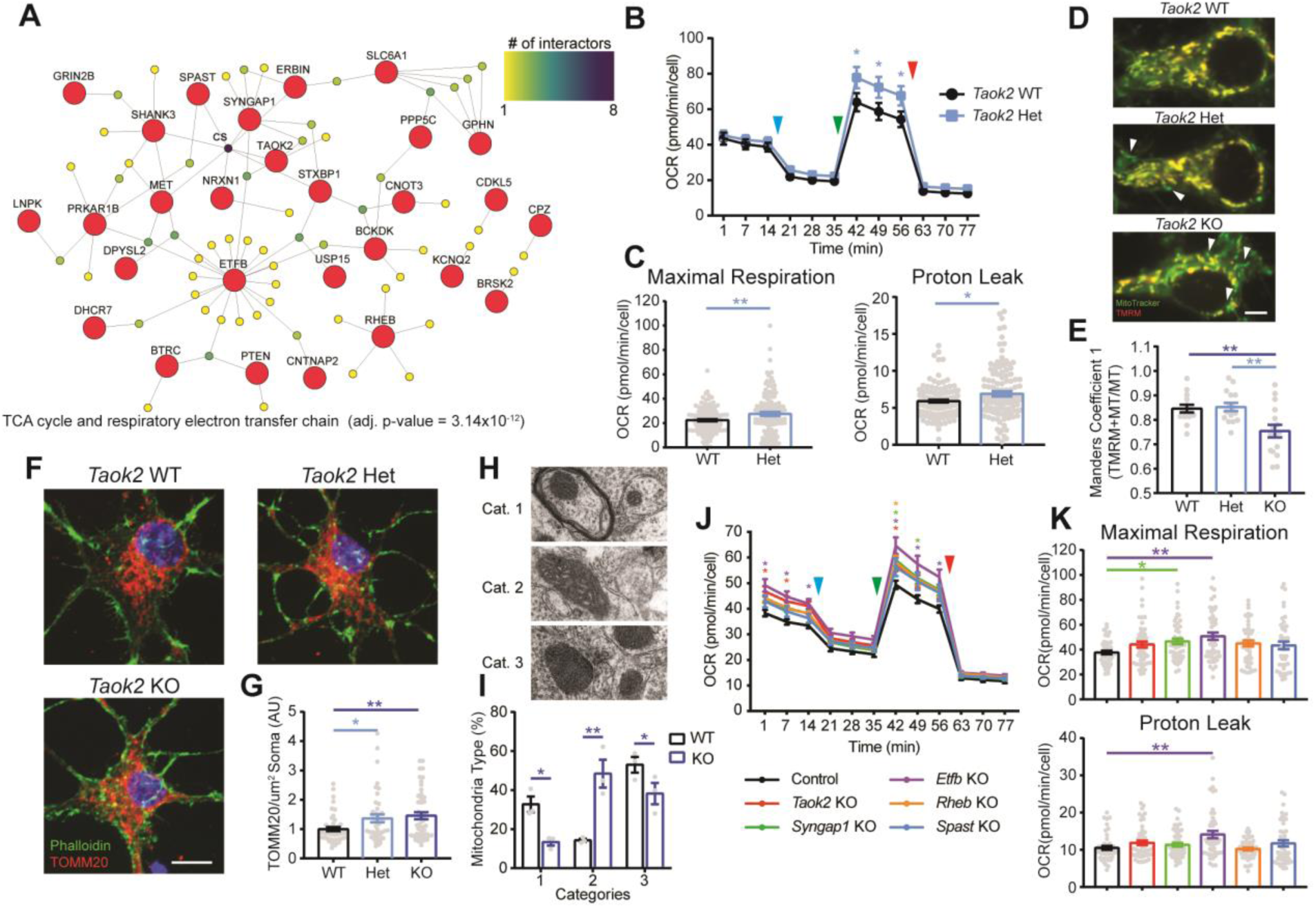
Regulation of cellular respiration and mitochondrial activity by ASD-risk genes. (A) Network of TCA cycle and pyruvate metabolism proteins enriched in the shared ASD-risk gene PPI network (g:Profiler, Benjamini-Hochberg FDR adj. p<0.05). (B) Loss of Taok2 alters cellular respiration. WT = 118 wells and Het = 122 wells from three separate cultures. Arrows indicate the addition of oligomycin (blue), FCCP (green), and Antimycin A and Rotenone (red). (C) Loss of Taok2 increases maximal respiration (*left*) and proton leakage (*right*). WT = 118 wells, Het = 122 wells from three separate cultures. (D) Representative images of Taok2 WT, Het, and KO neurons stained with MitoTrack (green) and TMRM (red). Arrows indicate Mitotracker-labeled mitochondria with no TMRM staining. Scale bar is 5 µm. (E) Taok2 Het and KO neurons have decreased active (TMRM) Mitotracker-labeled mitochondria. WT = 14, Het = 15, KO = 15 neurons from 1-3 separate pups per genotype from one culture. (F) Representative images of Taok2 WT, Het, and KO neurons stained with TOMM20 (red) and Phalloidin (green). Scale bar 5 µm. (G) Taok2 Het and KO neurons have increased TOMM20 staining. WT = 45, Het = 42, KO = 48 neurons from 1-4 separate pups per genotype from two cultures. (H) Representative images of synaptic mitochondria morphological categories. (I), Taok2 KO neurons have increased proportion of category 2 mitochondria with enlarged cristae. 19-25 images per animal and three animals per genotype. (J) CRISPR/Cas9 KO of Taok2, Syngap1, Etfb, Rheb, and Spast alters cellular respiration. Control = 45 wells, Taok2 KO = 51 wells, Syngap1 KO = 50 wells, Etfb KO = 47 wells, Rheb KO = 48 wells, Spast KO = 45 wells from five separate cultures. Arrows indicate the addition of oligomycin (blue), FCCP (green), and Antimycin A and Rotenone (red). (K) Maximal respiration (*top*) and proton leakage (*bottom*) show an increase in Syngap1 and Etfb KO neurons. Mean ± s.e.m. *p<0.05, **p<0.01, ***p<0.001.

To determine if other ASD risk genes converging on the mitochondrial/metabolic network regulate cellular respiration, we used the CRISPR/Cas9 system to knock out *Syngap1*, *Spast*, and *Taok2*. We also targeted *Etfb* and *Rheb*, both ASD risk genes that localize to the mitochondrion or regulate neuronal mitochondrial function (Yang et al., 2021). Combined gRNAs against BFP and Luciferase were used as a negative control (Hart et al., 2015; Richardson et al., 2016), and we used and validated 1-3 gRNAs targeting each gene (Figure S8D and S8E). CRISPR/Cas9 knockout of Etfb showed increased basal and maximal respiration, proton leakage, and no change in ATP synthase-dependent cellular respiration (Figure 3J and 3K, and Figure S8F and S8G). Mouse neurons with CRISPR knockout of *Taok2*, *Syngap1*, and *Rheb* also showed changes in many aspects of cellular respiration (Figure 3J and 3K, and Figure S8F-S8I). CRISPR KO of *Spast* did not cause changes in cellular respiration; however, Spast could regulate other aspects of the TCA cycle and pyruvate metabolism. The increase in basal respiration in *Taok2*, *Syngap1*, and *Etfb* KO neurons (Figure S8F) may be indicative of cellular respiration not yet reaching homeostasis (Ruggiero et al., 2021). We further validated the BioID data by using CRISPR/Cas9 to knockout Ap2s1, Gria1, Ppp1r9b, and Ppp2r5d; genes which did not have mitochondrial or metabolic proteins in their PPI network. Seahorse assays on these genes showed no changes in cellular respiration (Figure S8J-S8L). These findings suggest that a subset of ASD risk genes regulate cellular respiration in neurons, and highlight the relevance of TCA cycle and pyruvate metabolism pathways in ASD.

### PPI networks identify differences in signaling between missense variants in ASD risk genes

Next, we hypothesized that PPI networks could be used to identify differences in pathogenic mechanisms of *de novo* ASD-linked missense variants, in particular variants of unknown significance (VUS). Missense variants have been suggested to impact protein stability and protein-protein interaction networks (Chen et al., 2020); however, these data were imputed from databases using primarily non-neuronal datasets. Due to the strong link between synaptic functional deficits and ASD pathophysiology, we chose two known synaptic genes (TAOK2β, the synapse-specific isoform of TAOK2, and GRIA1) and a less well-characterized risk gene with no known cellular localization (PPP2R5D) (Figure 4A-4C, Table S8, and Table S9).

**Figure 4.**
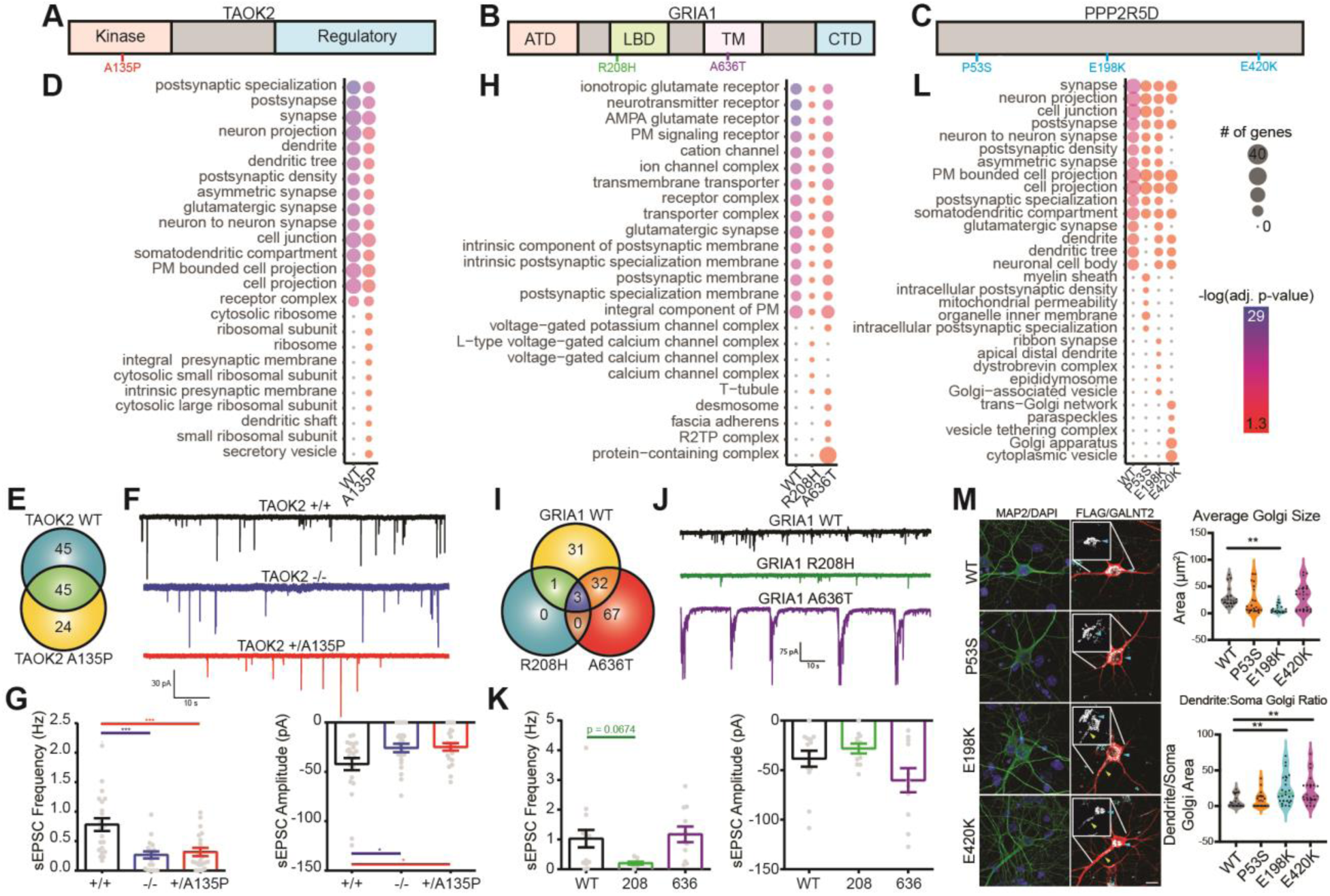
*De novo* missense variants alter the PPI networks of uncharacterized and established ASD-risk genes that correspond to functional deficits. (A-C), Diagram of TAOK2, GRIA1, and PPP2R5D and *de novo* missense variants. ATD: amino-terminal domain, LDB: ligand-binding domain, TM: transmembrane domain, CTD: carboxy-terminal domain, PM: plasma membrane. (D) Dot plot of top 15 cellular compartment gene sets and top 10 variant-specific gene sets in TAOK2 variants (g:Profiler, Benjamini-Hochberg FDR adj. p<0.05).. Size of dots indicate protein number and the color represents the significance. (E) Venn diagram of PPI network proteins of TAOK2 WT and A135P. (F) Representative traces of sEPSC recordings of DIV21 TAOK2 WT, KO, and A135P human iPSC-derived NGN2 neurons. (G) TAOK2 KO and A135P neurons show decreased sEPSC frequency (*left*) and amplitude (*right*). WT = 23, KO = 22, and A135P = 21 neurons from three transductions. (H) Dot plot of top 15 cellular compartment gene sets and top five variant-specific gene sets in GRIA1 variants (g:Profiler, Benjamini-Hochberg FDR adj. p<0.05). (I) Venn diagram of PPI network proteins of GRIA1 WT and variants. (J) Representative traces of sEPSC recordings of mouse neurons expressing GRIA1 or variants. (K) R208H variant shows trend in decrease sEPSC frequency (*left*) and no change in amplitude (*right*). WT = 14, R208H = 11, and A636T= 11 neurons from three transductions. (L) Dot plot of top 15 cellular compartment gene sets and top five variant-specific gene sets in PPP2R5D variants (g:Profiler, Benjamini-Hochberg FDR adj. p<0.05). (M) Representative images of PPP2R5D WT and variants show altered Golgi morphology and localization to the dendrite (white arrow) and soma (blue arrow) (*right*) through GALNT2 staining. Inlet shows magnified Golgi. Scale bar is 20µm. Neurons expressing the E198K variant have decreased Golgi size, while both the E198K and E420K variants cause mislocalization of the Golgi into the dendrite (*left*). 20 neurons from 5 separate infections per condition. Mean ± s.e.m. *p<0.05, **p<0.01, ***p<0.001.

We determined the change in the TAOK2β PPI network due to the A135P *de novo* missense variant. The TAOK2β A135P PPI network had reduced numbers of proteins associated with the synaptic compartment, and increased dendritic and ribosomal proteins (Figure 4D, Table S8 and Table S9). The latter changes may be due to the loss of PPIs in dendritic spines where TAOK2β localizes, causing an increase in dendritic and ribosome translation complex PPIs for the TAOK2β A135P (Figure 4E), combined with decreased expression of the A135P mutant (Figure S9A). To corroborate the possible synaptic deficits caused by the A135P variant, we performed patch-clamp electrophysiology on isogenic iPSC-derived NGN2-neurons (Figure S9B-S9C) (Deneault et al., 2018, 2019; Zhang et al., 2013). *TAOK2* KO and *TAOK2* A135P neurons had decreases in frequency and amplitude of spontaneous excitatory post-synaptic currents (sEPSCs) (Figure 4F and 4G). This data coincides with the reduced biotinylation of synaptic proteins for TAOK2 A135P-BioID2 (Figure 4D). The lack of change in the intrinsic firing properties or Synapsin1-positive punctae in *TAOK2* A135P neurons, as opposed to the *TAOK2* KO neurons (Figure S9B-S9E), suggest that the shift in interaction of proteins and localization for the heterozygous A135P line has dissimilar phenotypes compared to the *TAOK2* KO. In fact, *TAOK2* A135P neurons displayed increased size of Synapsin1 punctae, suggesting possible changes in synapse structure (Figure S9D and S9E). Taken together, these data demonstrate that changes in PPI networks can help to support functional deficits.

We also asked whether PPI networks can distinguish the impact of missense variants based on their location within functional domains of a gene. We investigated *GRIA1* and two ASD-linked *de novo* missense variants, R208H and A636T (Geisheker et al., 2017; Iossifov et al., 2014; de Ligt et al., 2012), located in the extra-cellular ligand binding domain and the transmembrane domain, respectively (Fig. 4B). The *GRIA1* variants showed differential effects in their enriched cellular compartments (Figure 4H) and the number of shared interacting proteins compared to wildtype (Figure 4I). GRIA1 R208H had a loss of proteins localizing to the AMPA receptor and post-synaptic density. GRIA1 A636T had a less severe impact, with small increases in compartment-specific PPIs, including membrane junction and ER proteins (Figure 4H, Table S8, and Table S9). There was no difference between GluA1 variant expression (Figure S9F) or in GluA2 subunit expression (Figure S9G). We identified electrophysiological changes that coincide with the changes in PPI networks, revealing a trend towards decreased sEPSC frequency in neurons expressing the R208H variant, but not the A636T variant (Figure 4J and 4K). Although the A636T mutant had no change in sEPSCs, we did observe large sEPSC bursts (Figure 4J), which may be indicative of altered trafficking of AMPA receptors (Pick and Ziff, 2018; Schwenk et al., 2019). These data demonstrate the potential use of PPI networks to identify functional differences in missense variants for receptor proteins.

Finally, we used BioID2 to test missense variants in the ASD risk gene PPP2R5D, a regulatory subunit of phosphatase-2A (Shang et al., 2016). We selected three *de novo* variants, P53S, E198K, and E420K (Houge et al., 2015; Shang et al., 2016). The PPI networks for the variants had both common and dissimilar effects (Figure 4C, 4L, and S9H), with all three variants reducing PPIs with synaptic and dendritic proteins (Figure 4L). All of the variants caused a loss and gain of diverse interactions (Figure S9H), with no change in expression levels (Figure S9I). Interestingly, both the E198K and the E420K variants gained trans-Golgi compartment proteins (Figure 4L, Table S8, and Table S9), suggesting altered localization to the Golgi apparatus. Previous studies have linked overactive AKT signaling to the PPP2R5D E420K variant (Papke et al., 2021); however, we found no difference in phospho-AKT levels (Figure S9J). To probe the E198K and E420K variants, we stained infected mouse neurons for a trans-Golgi apparatus-specific protein, GALNT2 (Figure 4M). In WT neurons, the GALTN2 expression is primarily located in the cell body (Figure 4M). However, GALTN2 staining in the E198K variant was smaller and spread throughout the primary dendrite (Figure 4L and 4M, and Table S8 and S9). Neurons expressing the E410K variants showed no change in size; however, GALTN2 expression was also spread into the apical dendrites (Figure 4L and 4M, and Table S8 and S9). As a control, we measured cell size and found that overexpression of PPP2R5D variants had no effect (Figure S9K). These data suggest that ASD-linked PPP2R5D variants may impart pathogenicity by altering Golgi-related functions in neurons.

### The 41 ASD-risk gene PPI network map enriches for additional ASD risk genes, human disease cell types, and correlates with human behavioral phenotypes from clinical datasets

To demonstrate the utility of the 41 ASD-risk gene PPI network map resource, we used enrichment analysis to determine relevance to human ASD. We found a significant enrichment of 112 additional ASD-risk genes (Fisher’s Exact test p = 2.69×10^-30^, OR = 3.45), highlighting the functional connectivity between ASD-risk genes at the protein level (Figure 5A). We also found that gene lists from individual sequencing studies were enriched in our PPI network, especially when examining cytoplasmic (non-nuclear) proteins (Figure S10A). Gene lists with only nuclear proteins were not enriched (Figure S10B), providing evidence for less interaction between proteins in the nucleus and the cytoplasm. Of the 153 ASD-risk proteins in the network, 69 are interacting with 2 or more ASD bait proteins.

**Figure 5.**
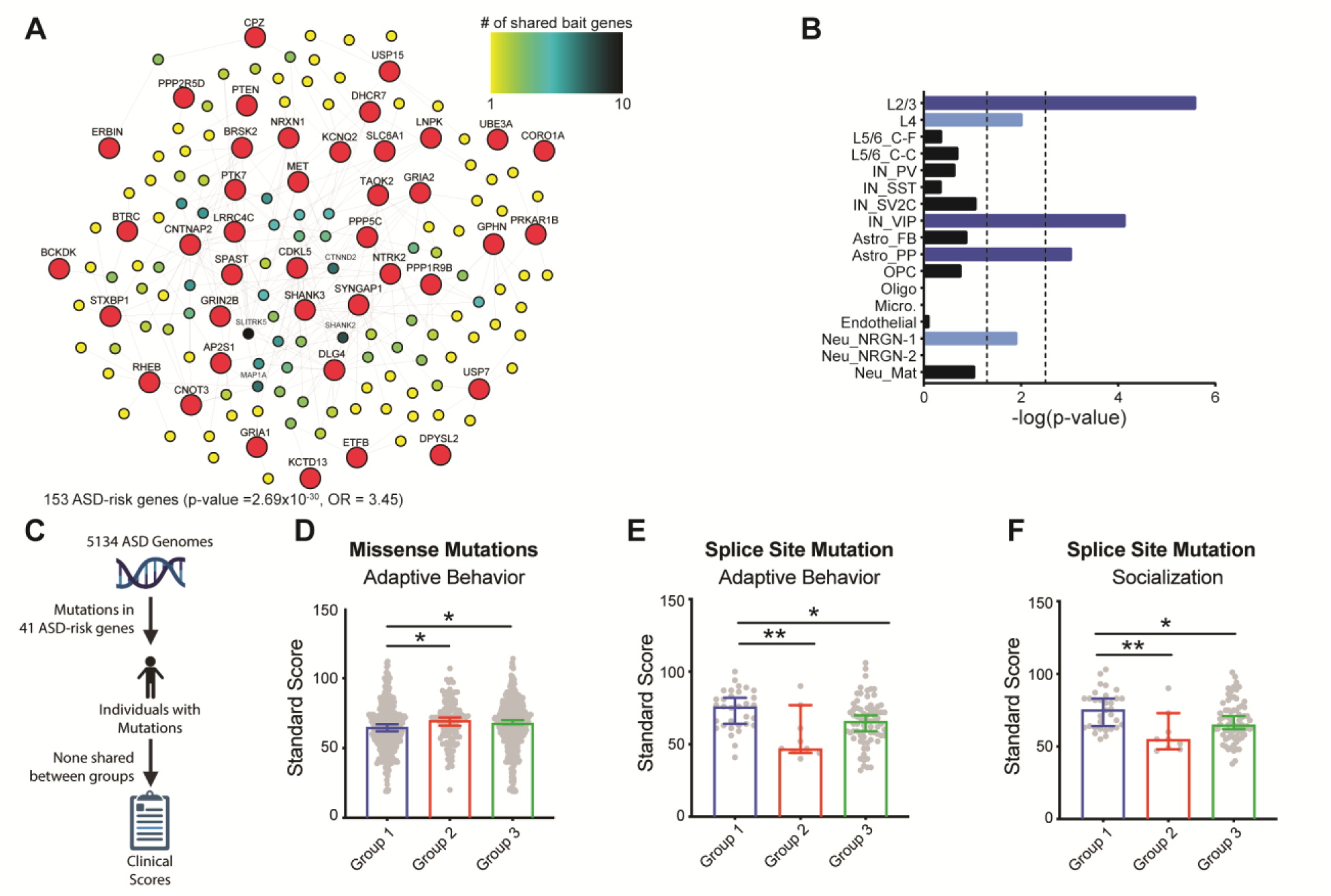
Shared ASD PPI network map network correlates to human brain development and disease pathology. (A) Network of ASD genes enriched in the shared ASD-risk gene PPI network. Large red nodes represent bait genes, smaller colored nodes represent sharedness of ASD-risk genes between bait genes, between 1 (yellow) and 10 (blue) shared bait genes (g:Profiler, Benjamini-Hochberg FDR adj. p<0.05). (B) Enrichment of ASD differentially expressed genes (DEGs) in cell types. (Fisher’s exact test). Dashed lines represent nominal (p = 0.05, *left*) and Bonferroni corrected (p = 0.05/number of cell types, *right*) significance thresholds. EN = excitatory neurons, IN = inhibitory neurons, RGC = radial glial cells, MGE RGC = medial ganglionic eminence, IPC = intermediate progenitor cells, Astro. = astrocyte, OPC = oligodendrocyte progenitor cells, Micro. = microglia, Endo. = endothelial cells, CP = choroid plexus cells, C-F = cortico-fugal, C-C = cortico-cortico, PV = paravalbumin, SST = somatostatin, VIP = vasoactive intestinal peptide, FB = fibrous, PP = protoplasmic, Neu_NRGN = neurogranin-expressing. Light blue bars have nominal p-value significance, while dark blue bars have Bonferroni corrected significance. (C) Flow chart of clinical data extraction (D) Decrease in the average standard scores of individuals with ASD, who have rare inherited missense mutations in Group 1 genes compared to Groups 2 and 3 (Group 1 = 350, Group 2 = 113, and Group 3 = 416 probands). Individuals with splice site mutations in Cluster 1 have significantly higher adaptive behavior (E) and socialization standard (F) scores than Cluster 2 and 3. Group 1 = 32, Group 2 = 9, and Group 3 = 71 probands. Mean ± s.e.m. *p<0.05, **p<0.01.

While the PPI network from 41 ASD-risk genes was generated using human genes, it was obtained using mouse cortical neurons; therefore, it is unknown whether this network map is applicable to human brain cell types or differentially expressed genes (DEGs) implicated in ASD pathology. We examined the enrichment of specific cell types based on their single cell RNA-sequencing profiles (Nowakowski et al., 2017; Velmeshev et al., 2019). We found the 41 ASD-risk gene PPI network map enriches for excitatory and inhibitory neuron cell types, and neural progenitor cells, astrocytes and microglia (Figure S10C), which have been associated with ASD (Parikshak et al., 2013; Tang et al., 2014; Velmeshev et al., 2019; Willsey et al., 2013; Xu et al., 2020). When examining the ASD-specific DEGs of cell types from human post-mortem brains (Velmeshev et al., 2019), the shared PPI network was enriched for DEGs in layer 2/3 and 4 neurons, parvalbumin and VIP interneurons, and protoplasmic astrocytes (Figure 5B). The enrichment of ASD DEGs of specific cell types suggests the human disease relevance of the 41 ASD-risk gene PPI network map.

Finally, we hypothesized a potential relationship between highly connected genes and human ASD behavioral phenotypes. We obtained clinical data of individuals with rare variants in the 41 ASD-risk genes from the MSSNG database and grouped them based on the correlation of individual ASD-risk gene PPI networks (Figure 2B and 5C) (Yuen et al., 2017). We obtained the adaptive behavior and socialization scores from between 112-879 individuals, which was dependant on data availability, and the number of individuals per group with at least one rare missense/splicing/LoF variant in the 41 ASD-risk genes (data-explorer.mss.ng). Remarkably, we found that individuals with missense variants in Group 1 genes had lower adaptive behavior standard scores compared to Groups 2 or 3, suggesting that missense variants impact Group 1 genes in regards to adaptive behavior (Figure 5D and Figure S10D). However, individuals with variants impacting mRNA splicing in Groups 1 had significantly higher standard adaptive behavior and socialization scores compared to Group 2 or 3 (Figure 5E and 5F). Interestingly, the NRXN1 gene that is part of group 2 has been found to have alternative splicing in individuals with neuropsychiatric disorders (Flaherty et al., 2019). No significant differences were seen between individuals with frame shift or stop gain variants (Figure S10E and S10F). Group 1 genes were found to have the largest enrichment of ASD-risk genes (Figure S10G), suggesting that the highly interconnected PPI networks and shared pathways for this group of genes may be linked to the clinical phenotypes.

### Discussion and Conclusion

A long-standing question in ASD research is whether there are convergent signaling mechanisms between different risk genes. Specific disease cell types or signaling pathways have been proposed as convergent mechanisms, but these data are based on RNA expression, not protein levels (Chang et al., 2015; O’Roak et al., 2012b; Ramaswami et al., 2020; Velmeshev et al., 2019; Voineagu et al., 2011). To address this, we devised a neuron-specific proteomic screen to study 41 ASD-risk genes, and identified links between risk genes and convergent pathways. In addition, PPI network mapping could predict the impact of disease-associated missense variants. Finally, the PPI network map revealed an enrichment of additional ASD risk genes and cell types implicated in ASD pathology. Cross-referencing the PPI network with human clinical data showed correlation between highly interacting ASD-risk genes and ASD diagnostic behavioral severity.

A main finding was the identification of shared interactions with the TCA cycle and mitochondrial proteins in 28 ASD-risk genes, where dysfunction is indirectly associated with neurodevelopmental disorders (Iwata et al., 2020) (Figure 3A). Clinical studies have found mitochondrial and metabolic dysfunction or changes in metabolites in primary lymphocytes or brain tissue in individuals with ASD, but whether this is direct or indirect is not known (Anitha et al., 2012; Frye and Rossignol, 2011; Frye et al., 2013; Kurochkin et al., 2019; Rose et al., 2014, 2018; Rossignol and Frye, 2012; Wang et al., 2016b). Some ASD-associated syndromic disorders, co-morbid disorders and genetic ASD models have shown deficits in mitochondrial and metabolic processes, however the specific proteins involved were unknown (Bülow et al., 2021; Ebrahimi-fakhari et al., 2016; Fernandez et al., 2019; Jagtap et al., 2019; Kanellopoulos et al., 2020; Li et al., 2019; Licznerski et al., 2020; Madison et al., 2021; Menzies et al., 2021; Shen et al., 2019b; Shulyakova et al., 2017). Our findings indicate that TCA cycle and mitochondrial activity proteins are interacting with multiple ASD-risk genes. In addition, we identified two ASD-risk genes (*RHEB* and *BCKDK*) that were previously implicated in metabolic processes (Heinemann-Yerushalmi et al., 2021; Yang et al., 2021). This highlights that our screen can identify relevant protein interactions and may indicate a more direct connection between mitochondrial/metabolic processes and some ASD genes (Jagtap et al., 2019; Madison et al., 2021).

Our CRISPR/Cas9 KO studies revealed that multiple ASD-risk genes are important for proper cellular respiration (Figure 3A), suggesting that cross-talk may exist between ASD-risk genes and TCA cycle function. Deficits in the TCA cycle can cause overreliance on glutaminolysis to produce energy and cause a decrease in synaptic vesicle glutamate levels, possibly explaining the synaptic transmission deficits caused by *Syngap1* and *Taok2* KO (Divakaruni et al., 2018; Fendt and Verstreken, 2017; Namba et al., 2021).

ASD-associated *de novo* missense mutations are enriched in hub genes of known protein interaction networks (Chen et al., 2018, 2020). However, few studies have used proximity-labeling to study disease-relevant mutations on PPI networks (Chou et al., 2018; Pintacuda et al., 2021; Tracy et al., 2021). Our screen provides functional evidence of the impact of *de novo* missense variants on the PPI network of three ASD-risk genes as examples. PPP2R5D is the regulatory subunit of PP2A, which was previously associated with Golgi assembly through the beta subunit (Lowe et al., 2000; Schmitz et al., 2010). We found PPP2R5D variants disrupted the Golgi apparatus morphology and localization, highlighting a potential pathogenic mechanism in neurons. These studies are an example of how PPI networks can be used to discover unknown disease mechanisms. Using neuron-specific PPI networks allows the use of a relevant cell type, and the potential to study variants of unknown significance to provide important information on severity.

The enrichment of 112 ASD risk genes in the shared ASD-risk gene PPI network map, and the enrichment of the network in ASD-associated cell types further emphasizes the interconnectedness of ASD risk proteins. Mid-fetal deep cortical projection neurons and superficial cortical glutamatergic neurons are enriched for ASD-risk genes and are associated with autism pathology (Parikshak et al., 2013; Willsey et al., 2013). The ASD-shared PPI network was highly enriched for genes expressed in excitatory and inhibitory neurons, and for DEGs in individuals with ASD specific to layers 2/3 excitatory neurons and VIP interneurons. Future studies will need to distinguish which ASD PPI networks are specific to each cell type, or possible subpopulations, to understand the subtle network changes that impact disease mechanisms.

The grouping of the 41 ASD-risk genes based on their PPI network, and correlation to clinical behavioral scores is an emerging area. Although we focused on missense/LoF variants, we found that the type of variant in each group of genes influenced the average score of the individuals within the group. However, the overall effect size of the analysis was small, suggesting that further BioID of ASD-risk genes, or methods that consider the genetic background of individuals, is required. To work through the complexity, it may be important to combine our analysis with other methods of categorizing mutations (e.g., gnomAD pLI, Polyphen-2) as higher or lower impact, which would reduce the number of people shared between groups (Adzhubei et al., 2010; Lek et al., 2016).

In conclusion, our neuron-specific 41 ASD-risk gene PPI network map demonstrates that interacting and close proximity proteins, and their associated pathways and sub-cellular compartments are relevant to ASD disease pathology, which is missing from transcriptome-based approaches. This resource containing the individual PPI networks of 41 ASD-risk genes will be valuable for future in-depth study of the genes, and has the potential to grow larger with PPI networks of additional risk genes.

### Limitations of the study

In our screen, we excluded nuclear genes due to the functional separation with cytoplasmic genes (Satterstrom et al., 2020). Therefore, we did not identify many disease-relevant convergent pathways associated with gene regulation and chromatin modification. Another limitation is the variability between BioID runs that necessitated reduced stringency in our comparisons. Therefore, functional validation is important to filter out changes caused by background noise. Finally, we used single canonical isoforms and C-terminal tagged BioID (no N-terminal tag) of each gene; therefore some PPI networks may not encompass the full scope of possible interactions in the neuron.

## Acknowledgements

We thank Courtney Irwin. and Paolo de Guzman for proof reading the manuscript. We also thank Kristin J. Brennand and Michael B. Fernandez for providing the NRXN1 cDNA. Graphical abstract and flowcharts were created with BioRender.com (BU24JATEEX, SD235B8ORF, KW235KT7TM, RZ235KTA0S).

## Author contributions

N.M. and K.K.S. conceived the project. N.M. and K.K.S. wrote the paper with input from A.A.C., B.T., W.E., B.T., and B.W.D. A.A.C and B.K.U created the BioID2 lentiviral construct. N.M. generated all subsequent DNA constructs and performed all experiments and data analysis unless otherwise specified. N.M and A.A.C. generated all lentiviruses. S.X. and Y.L. ran mass spectrometer samples and data acquisition. C.O.B. performed electrophysiology recordings. J.A.U, J.E.H., and N.P. assisted with western blots. D.P.M., S.H., B.S., and F.C.dA. performed and analysed mitochondrial experiments. E.D., J.E, and S.W.S helped to create the human *TAOK2* KO and A135P iPSC lines. E.A. advised on clinical score analysis and G.D.B. advised on pathway analyses. K.S.S supervised the project.

## Inclusion and Diversity

We support inclusive, diverse, and equitable conduct of research.

## RESOURCE AVAILABILITY

### Lead contact

Further information and requests for resources and reagents should be directed to and will be fulfilled by the Lead Contact, Dr. Karun K. Singh

### Material availability

All constructs and material are available by request from Dr. Karun K. Singh.

### Data and code availability

- Mass spectrometry datasets consisting of raw files and results files with statistical analysis to identify PPI networks are available via ProteomeXchange with identifier PXD036946. DEPs are available via ProteomeXchange with identifier PXD036937. Individual PPI networks and shared ASD-risk gene PPI network map protein lists and enriched pathways can be found in Tables S1-S9. The Mouse_Human_Reactome and Mouse_GO_ALL_no_GO_iea gene sets used for overrepresentation and gene set enrichment analyses were downloaded on 13 August 2021 from http://download.baderlab.org/EM_Genesets/(Merico et al., 2010). RNA sequencing raw sequence files and results files with statistical analysis to identify significant DEGs are available through the Gene Expression Omnibus (Accession: GSE213899). ASD proband variant information and clinical scores are available through the MSSNG database (research.mss.ng)(Yuen et al., 2017) and the associated Metabase (data-explorer.mss.ng), respectively.
- This paper does not report original code.
- Any additional information required to reanalyze the data reported in this work paper is available from the Lead Contact upon request.

## METHOD DETAILS

### Antibodies

The following antibodies were used for immunostaining and immunoblotting experiments: rabbit anti-FLAG (IB 1:2,000, MilliporeSigma, F7425), mouse anti-FLAG (IF 1:1,000, IB: 1:2000, MilliporeSigma, F3165), rabbit anti-turboGFP (IF 1:1,000, IB 1:1,000, Fisher, PA5-22688), chicken anti-MAP2 (IF 1:1,000, Cedarlane, CLN182), rabbit anti-β-actin (IB 1:1,000, Cell Signaling, 8457S), mouse anti-β-actin (IB 1:5,000, MilliporeSigma, A5316), goat anti-TAOK2α/β (IB 1:1,000, Santa Cruz Biotechnology, sc-47447), rabbit anti-TAOK2β (IB 1:1,000, Synaptic Systems, 395 003), mouse anti-Synapsin1 (IF: 1:1000, Synaptic Systems, 106 001), mouse anti-TOMM20 (IF 1:100, US Biological, 134604), mouse anti-Shank2 (IF 1:500, Antibodies Inc), mouse anti-PSD95 (IF 1:1000, NeuroMab), rabbit anti-FMRP (IB 1:1000, Abcam), rabbit anti-STXBP1 (IB 1:1000, Proteintech), mouse anti-CS (IB 1:1000, Thermofisher), rabbit anti-CS (IP, Proteintech), rabbit anti-SYNGAP1 (IB 1:1000, Thermofisher). DAPI (IF 300mM, ThermoFisher, D21490), Hoechst (IF 1:10,000, Invitrogen, 1050083), Phalloidin-488 (IF 1:120, Cytoskeleton Inc., PHDG1), Anti-mouse-Cy3 (IF 1:500, Jackson Immunoresearch, 715-165-151), Alexa 488 anti-rabbit (IF 1:500, Jackson Immunoresearch, 711-545-152), Alexa 488 anti-chicken (IF 1:500, Jackson Immunoresearch, 703-545-155), 405 conjugated-streptavidin (IF 1:500, Jackson Immunoresearch, 016-470-084), 405 anti-chicken (IF 1:500, Jackson Immunoresearch, 703-475-155), Alexa 647 anti-mouse (IF 1:500, Jackson Immunoresearch, 715-605-150).

### Generation of constructs

All cloning was accomplished using the In-Fusion HD cloning kit (Takara). To create the BioID2 fusion constructs, we obtained an expression construct containing a 198bp (13x “GGGGS” repeat) linker sequence upstream of a C-terminal 3xFLAG-tagged BioID2 sequence with BioID2 (Genscript). For lentiviral expression, 13xlinker-BioID2-3xFLAG was amplified and cloned into the lentiviral backbone pLV-hSYN-RFP (Addgene #22909)(Nathanson et al., 2009). For ease of visualization and to create a bicistronic construct, the RFP in the pLV-hSYN-RFP backbone was replaced with the TurboGFP(tGFP)-P2A from pCW57-GFP-2A-MCS (Addgene #71783)(Barger et al., 2019). NheI digest sites were added after the P2A sequence and before the 13xLinker to allow easy insertion of ASD-risk bait genes. The final construct being pLV-hSyn-tGFP-P2A-Bait-13xLinker-BioID2-3xFLAG (referred to as the BioID2 fusion construct). For the control luciferase construct a second P2A was cloned in between the luciferase ORF and the 13xLinker, creating the pLV-hSyn-tGFP-P2A-Luciferase-P2A-13xLinker-BioID2-3xFLAG construct (referred to as the Luciferase control construct). ASD-risk genes open reading frames (ORFs) were purchased from Addgene and Genscript or amplified from human adult and fetal brain RNA (Takara) (see Table S10)(Alford et al., 2012; Braganza et al., 2017; Braun et al., 2011; Butko et al., 2012; Cummins and Vogelstein, 2004; Furlong et al., 2000; Hiday et al., 2017; Howarth et al., 2005; Johannessen et al., 2010; Kim et al., 2016; Lu et al., 2014; Małecki et al., 2015; Ohno et al., 2002; Seeling et al., 1999; Solowska et al., 2010; Sowa et al., 2009; Urano et al., 2007; Wang et al., 2016a). For mouse electrophysiology experiments, the GRIA1, GRIA1 R208H and GRIA1 A636T ORFs were inserted between the GFP-P2A and 3xFLAG. The pLV-CMV-Cas9-T2A-EGFP plasmid was made by replacing the UBC promoter-rTetR in the FUW-M2rtTA plasmid (Addgene #20342)(Hockemeyer et al., 2008) with CMV-Cas9-T2A-EGFP from PX458 (Addgene #48138)(Ran et al., 2013). All generated constructs are available upon request.

### Experimental Models

Taok2 Het (*Taok2* +/-) and KO (*Taok2* -/-) mice were created by Kapfhamer *et al*.(Kapfhamer et al., 2013). The E15-16 or E18 mouse embryo brains were used for cortical neuronal cultures. P21-P23 mice were used for mass spectrometry or RNA sequencing experiments. Animals housed at the Central Animal Facilities at McMaster University were approved for experiments and procedures by the Animal Research Ethics Board (AREB) at McMaster University. Animals housed at the University Medical Center Hamburg-Eppendorf, Hamburg were approved for experiments and procedures by local authorities of the city-state Hamburg (Behörde für Gesundheit und Verbraucherschutz, Fachbereich Veterinärwesen) and the animal care committee of the University Medical Center Hamburg-Eppendorf. All procedures were performed according to the German and European Animal Welfare Act. Animals housed at the Animal Resource Center at University Health Network were approved for experiments and procedures by the University Health Network animal care committees.

All work with the human iPSCs was performed with the approval of the Hamilton Integrated Research Ethics Board. iPSCs were previously generated as described in (Deneault et al., 2018).

### Subject Details

No subjects were directly tested in this study. Adaptive behavior and socialization standard scores of affected individuals was extracted from the MSSNG associated Metabase (data-explorer.mss.ng). These individuals are part of the MSSNG database (research.mss.ng)(Yuen et al., 2017), which has whole genome sequences of 4,258 families and 5,102 ASD-affected individuals.

### Mouse Cortical Neuron Cultures

E15-E16 CD1 mice (Charles River) embryo cortices were harvested using a dissecting microscope and kept in HBSS. Cortices were then digested in 300 µg/ml of papain (Worthington) and 2 U/ml of DNase (Thermo) for 20 minutes at 37 °C. Cortices were then washed three times with mouse plating media (Neurobasal media supplemented with 2 mM GlutaMAX (Thermo), Pen-Strep (Thermo), and 10% FBS(Gibco)). Digested cortices were triturated and put through 40 µm strainer. Cells were counted, suspended in plating media, and plated at 600,000 cells per well of a 12-well plate. Plates were coated with 100 µg/ml poly-D-lysine (mol wt > 300,000, Sigma) and 3 µg/ml Laminin (Sigma). For immunostaining, 12 mm coverslips (Fisher) were placed in the well prior to coating. The cells were incubated at 37 °C (with 5 % CO_2_) for one hour, after which plating media was removed and replaced with mouse culturing media (Neurobasal media supplemented with 2 mM GlutaMAX, Pen-Strep, and B27). Cells were grown at 37 °C (with 5 % CO_2_) and half media changes were done on day 7 and every 3-4 days onwards.

### CRISPR/Cas9 editing of human induced pluripotent stem cells (iPSCs)

Human iPSCs were maintained on Matrigel (Corning) coated plates using mTeSR1 media (Stem Cell Technologies) and passaged every 3-4 days using ReLeSR (Stem Cell Technologies). Human iPSCs were edited for homozygous knockout of *TAOK2* or heterozygous knock-in of the A135P mutation as described in Deneault *et al*.(Deneault et al., 2018). MGB probes were ordered from ThermoFisher scientific and ssODN were designed on Benchling.com (Biology Software) and ordered from Integrated DNA Technologies. For the A135P mutation a mutant and wildtype ssODN containing the A135P (G to C) mutation and a PAM site mutation or just the PAM site mutation, respectively, were used to create a heterozygous knock-in.

### Human iPSC to neuron differentiation via NGN2 induction

Human iPSCs were cultured on Matrigel (Corning) coated plates using mTeSR1 media (Stem Cell Technologies) and passaged every 3-4 days using ReLeSR (Stem Cell Technologies) until neural induction. A modified NGN2 induction protocol (Zhang *et al*. 2013) was used to differentiate human iPSCs into excitatory NGN2 neurons(Zhang et al., 2013). Human iPSCs were dual infected with pTet-O-NGN2-P2A-EGFP and FUW-M2rtTA lentiviruses for dox-inducible expression and were titered for > 90% infection efficiency. On Day -1 iPSCs were singularized using Accutase (Stem Cell Technologies) and plated with mTeSR1 media (supplemented with 10 µM Y-27632) on Matrigel at 400,000 cells per well in a 6-well plate. On Day 0, media exchanged and supplemented with Doxycycline (1 µg/ml). On Day 1 and 2, media was replaced with iNPC media (DMEM/F12 media (Gibco) supplemented with N2 (Gibco), MEM NEAA (Thermo), 2mM GlutaMAX, and Pen-Strep) with Doxycycline and Puromycin (2µg/ml). On Day 3, media was then replaced with iNi media (Neurobasal media with SM1 (Stem Cell Technologies), 2mM GlutaMAX, Pen-Strep, 20 ng/ml BDNF, 20 ng/ml GDNF, and 1 µg/ml Laminin) with Doxycycline. On day 4, differentiated neurons were singularized using Accutase and re-plated at 100,000 cells per well in a 24-well plate in only iNi media. Plates were pre-coated with 20 µg/ml Laminin and 67 ug/ml Poly-ornithine (Sigma). Mouse glial cells were plated on top of the differentiated neurons after 24 hours at a density of 50,000 cells per well. Half-media changes were carried out every other day, and iNi media was supplemented with 2.5 % FBS on Day 9 and onwards. Neurons were grown until day 28 post NGN2-induction.

### Generation of high-titer lentivirus

All viruses were made using the 2^nd^ generation lentiviral packaging systems in Lenti-X HEK293 FTT cells (Takara). Lenti-X cells were passaged maximum 3 times before being used for virus production in HEK media (High glucose DMEM with 4 mM GlutaMAX, 1 mM Sodium Pyruvate, and 10 % FBS). Lenti-X cells were passaged once with 500µg/ml Gentamycin (Thermo) to increase T antigen expressing cells. Cells were plated into T150 flasks and each flask was transfected with the BioID2 lentiviral plasmid and the packaging plasmids, pMD2.G and pPAX2 (Addgene #12259 and #12260), using Lipofectamine 2000 in a 3:5 Opti-MEM: HEK media mix. Media was exchanged for fresh media after 5.5-6 hours. Media was harvested twice, first at 48 hours and then at 72 hours post-transection and spun at 100,000xG for 2 hours (maximum acceleration and deceleration). The virus was resuspended in PBS and kept at −80°C until they were used. Larger and unstable viruses were spun at 20,000xG for 4 hours in a table top centrifuge using a 20 % sucrose cushion(Yacoub et al., 2012). See nature exchange protocol for detailed procedure.

### Infection of mouse cortical neurons for BioID2 screen

One 12-well plate of 7.2 million mouse cortical neurons was considered as one biological replicate. Each cortical neuron culture produced at least 5 plates for four separate BioID2 bait gene samples and one luciferase control sample. Three separate cultures were done in a 3 days span in one week to get 3 biological replicates per protein-of-interest (POI). On days *in vitro* (DIV) 14, the conditioned media from the mouse neuron cultures were removed, leaving only 0.5 ml of media per well. Extra wells with and without coverslips were infected at the same MOI for flow cytometry measurements of GFP positive neurons and immunostaining, respectively. On DIV14, lentivirus with BioID2 fusion constructs were added to each well with an average transduction efficiency of 70%, and on DIV17 each well was supplemented with 50 µM of Biotin. After 18-20 hours, cells for mass spectrometry were lysed with RIPA buffer (1 % NP40, 50 mM Tris-HCl, 150 mM NaCl, 0.1 % SDS, 0.5 % deoxycholic acid, and protease inhibitor cocktail (PIC)) and flash frozen in liquid nitrogen. Cells for flow cytometry were dissociated with 0.25 % Trypsin-EDTA (Fisher) and resuspended in PEF media (PBS with 2 mM EDTA and 5 % FBS) (See flow cytometry section). Cells for immunostaining were fixed with 4 % PFA for 20 minutes, washed with PBS, and kept at 4 °C for staining.

### Transfection of HEK293 FT cells for BioID2 screen

10 million HEK293 FT cells were plated in a 10 cm culture dish and transfected 24 hours later with the BioID2 fusion construct plasmids using Lipofectamine 2000. Media was changed 6 hours after transfection and 50µM biotin was added 48 hours post-transfection. Cells were lysed 72 hours post-transfection in RIPA buffer and flash frozen in liquid nitrogen. Each individual plate was considered as biological replicate and three plates were used for each gene and the luciferase control. An extra plate was used for flow cytometry measurements of GFP positive cells.

### Processing of mouse cortical neuron and HEK293 FT cell BioID2 samples

Lysed cells were thawed and DNA was digested using benzonase (Sigma). Lysates were than sonicated at high speed for 5 seconds and centrifuged at 20,000xG for 30 minutes. The lysate supernatants were incubated with streptavidin Sepharose beads (GE Healthcare) at 4 °C for 3 hours. Following the incubation, the supernatant was spun down at 100xG for 2 minutes and the supernatant was removed. The beads were than washed once with RIPA buffer, and then six times with 100 mM triethylammonium bicarbonate (TEAB) with centrifugation between each wash. After the final wash, the beads are then resuspended in 100 mM TEAB and sequencing-grade trypsin (Promega) was added to digest the biotinylated proteins on the beads into peptides. The beads were incubated at 37 °C for 16 hours while rotating, and additional trypsin was added and incubated for a further 2 hours. The beads were then pelleted and the supernatant was transferred to a new tube. The beads were washed twice with 100 mM TEAB and each wash was added to the supernatant. The supernatant was then transferred to a 1.5 mL screw cap tub and speed vacuum dried. The dried peptides were stored at 4 °C for TMT-labeling.

### Multiplex TMT-labeling of BioID2 samples

TMT10plex isobaric-labeling was used to combine at least 3 biological replicates per gene. One additional technical TMT-labeling replicate of luciferase control sample chosen at random was used to account for differences in labeling. Dried peptides were resuspended in 100 mM TEAB. Each sample was TMT-labeled using the TMT 10plex Isobaric Mass Tagging Kit (Thermo). The four genes (proteins-of-interest, POI) were divided into two separate batches and the luciferase control samples were divided between the batches. Each batch had three biological replicates of the two genes and the luciferase control. One luciferase sample chosen at random was divided and labeled with two different labels to determine variance due to labeling efficiencies. In brief, TMT-label resuspended in acetonitrile was added to each sample and incubated at room temperature for one hour. To stop the reaction, 5 % hydroxylamine was then added to the samples and incubated for 15 minutes at room temperature. All ten samples were combined into one tube and divided into two samples. Both samples were then speed vacuum dried. One sample was kept at −80 °C for storage and the second sample was kept at 4 °C to be run in the mass spectrometer.

### Identification of biotinylated proteins from BioID2 screen samples using LC-MS/MS

Peptide samples were resuspended in 0.1% Trifluoroacetic acid (TFA) and loaded for liquid chromatography, which was conducted using a home-made trap-column (5 cm x 200 µm inner diameter; POROS 10 µm 10R2 C18 resin) and a home-made analytical column (50 cm x 50 µm inner diameter; Monitor 5 µm 100A C18 resin), running a 120min (label free) or 180min (TMT) reversed-phase gradient at 70nl/min on a Thermo Fisher Ultimate 3000 RSLCNano UPLC system coupled to a Thermo QExactive HF quadrupole-Orbitrap mass spectrometer. A parent ion scan was performed using a resolving power of 120,000 and then up to the 20 most intense peaks were selected for MS/MS (minimum ion count of 1000 for activation), using higher energy collision induced dissociation (HCD) fragmentation. Dynamic exclusion was activated such that MS/MS of the same m/z (within a range of 10 ppm; exclusion list size = 500) detected twice within 5 seconds were excluded from analysis for 30 seconds. Data were analyzed using Proteome Discoverer 2.2 (Thermo). For protein identification, search was against the Swiss-Prot mouse proteome database (55,366 protein isoform entries)(Bateman et al., 2021), while the search parameters specified a parent ion mass tolerance of 10 ppm, and an MS/MS fragment ion tolerance of 0.02 Da, with up to two missed cleavages allowed for trypsin. Dynamic modification of +16@M was allowed.

### Analysis for the identification of ASD-risk and cellular compartment protein PPI networks

Only proteins identified with two unique peptides were used for analysis. Two statistical cut-offs were used to identify positive hits for the PPI networks of each POI: Biotinylated proteins in the POI sample with 1) a significant increase in Log2 abundance compared to the luciferase control (Student’s t-test, p<0.05)(Uezu et al., 2016) and 2) that were significant outliers when accounting for the overall protein abundance compared to the protein abundance ratio between the POI and control samples (SigB p<0.05)(Cox and Mann, 2008). Protein abundances were normalized between biological replicates based on the sample with the highest total protein abundance. To reduce variability between each viral transduction, flow cytometry was used to determine the total abundance of GFP in the infected neurons between samples. The protein abundance levels of samples that had less total GFP (area under the curve in GFP intensity histogram) than the luciferase control were normalized by the factor required to equalize the total GFP of the POI sample to the luciferase control sample. To account for false positive hits due to variability in TMT-labeling between samples, the ratio of protein abundances between the luciferase control technical TMT-labeling replicates were used as the minimal required ratio between POI and control sample abundances. Proteins that did not have abundance ratios (POI/Luciferase control) higher than this minimal ratio were considered false positives and removed from further analysis.

### Pathway enrichment analyses

All pathway enrichment analysis was done using the g:Profiler GOst functional profiling tool (https://biit.cs.ut.ee/gprofiler/gost)(Raudvere et al., 2019). We used internal sources without electronic GO annotations for GO biological processes and GO cellular component (compartment), and curated Reactome pathway gene sets from the Bader lab (http://download.baderlab.org/EM_Genesets/)(Merico et al., 2010). All three sources were used for the shared ASD-risk gene network proteins. Only GO cellular component enrichment was used for the HEK293 FT cell BioID2 PPI networks, neuron cellular compartment BioID2 PPI networks, and *de novo* missense mutation network BioID2 PPI network comparisons. We compared the protein lists against a custom statistical domain of proteins identified through fractionated mass spectrometry of the mouse brain(Sharma et al., 2015) and combined with any additional proteins identified in the BioID2 screen. The final mouse brain proteome background had a total of 11992 proteins after removing multiple isoforms of the same protein. HEK293 BioID2 PPI networks were compared to all annotated gene lists. The g:Profiler Benjamini-Hochberg FDR multiple correction was used and only pathways with an adj. p-value < 0.05 were considered significantly enriched. For cellular component enrichment for *de novo* missense variant BioID2 PPI networks, the ggplot package in R was used to create the dot plots. For *de novo* missense variant BioID2 bait genes, all proteins identified in the wildtype samples were used for analysis, while for the shared PPI network map only proteins found in all wildtype samples were used for pathway enrichment analysis.

### Virus titering and GFP normalization for BioID2 screen

Mouse cortical neurons were cultured as described above and infected on DIV3 at three dilutions of virus (1:100, 1:333, 1:1000). On DIV 5, infected mouse cortical neurons were singularized using 0.25 % Trypsin-EDTA (Fisher) and resuspended in PEF media (PBS with 2 mM EDTA and 5 % FBS). For GFP normalization, DIV18 mouse neurons infected with the BioID2 lentiviruses were dissociated with Trypsin and resuspended in PEF media. CytoFLEX-LX or CANTO II flow cytometers (UHN Research Flow Cytometry Facility – KDT Site) were used to measure the percentage of GFP-positive cells with the 488 laser and 525/40 or 525/50 filters, respectively, using CytExpert software (Beckman Coulter). Functional titers were calculated based on the linear relationship between virus amount and percent of GFP positive cells. Mouse cortical neurons were infected at an MOI of 0.7, where 70 percent of cells were expected to be infected with the BioID2 lentiviral constructs. For normalization, the total GFP per 20,000 GFP-positive cells were quantified by taking the area under the GFP intensity histogram. GFP percentage and total amount was calculated using FlowJo software.

### Western blots

HEK293 FT cells were transfected with the BioID2 constructs using Lipofectamine 2000 (Invitrogen) in Opti-MEM: HEK media. Cells were harvested 48 hours post-transfection and lysed with RIPA buffer (with fresh PIC). Lysates were either snap-frozen in liquid nitrogen or taken directly for western blot sample preparation. Thawed or fresh lysates were sonicated at high frequency for 5 seconds and centrifuged at 20,000xG for 5 minutes at 4 °C. Lysates were than quantified using the Bio-Rad Bradford protein assay (Bio-Rad) by measuring absorbance with the SPECTROstar Nano machine and MARS Data analysis software (BMG LABTECH) and diluted to equal concentrations with RIPA buffer. 30-50 µg of protein were run on 8 % or 10 % SDS-PAGE Tris-Glycine gels (depending on the size of the proteins) at 100V for initial stacking and then 140V for 1-1.25 hours in a Tris-Glycine running buffer. Proteins were then transferred onto PVDF membrane using a Tris-Glycine buffered wet transfer system for 2 hours at constant 200 mA. Blots were then blocked with 5% milk in TBS-T (Tris buffered saline pH 7.4 with 0.1 % Tween). Blots were incubated with primary antibodies overnight in 5 % milk/TBS-T. The next day, membranes were washed three times with TBS-T for 5 minutes each and then incubated with secondary antibodies in 5 % milk/TBS-T for 1 hour. Blots were imaged by incubating them with the Amersham ECL western blotting detection reagent (VWR) for 1 minute and then imaging every 10 seconds for 5 minutes on the ChemiDoc XRS+ machine (Bio-Rad). ImageLab (Bio-Rad) was used for band intensity quantification.

### Co-immunoprecipitation Western Blots

Cortices were harvested from 3-4 week old CD1 mice and homogenized in NP40 lysis buffer (150 mM NaCl, 1% NP40, 50 mM Tris-Cl pH 8.0). Magentic dynabeads Protein G or Protein A (Invitrogen) were washed with 0.05% Triton X-100 in PBS, and then incubated with 4 µg of the antibody, or IgG for the control (Santa Cruz). Antibody-bead complexes were then washed 4 times with NT2 buffer (50 mM Tris-HCl, 150 mM NaCl, 1 mM MgCl_2_, 0.05% NP40). Lysates were quantified using the Bradford assay and 1 mg of protein was incubated with the antibody-bead complexes for 16hrs at 4 degrees Celsius. The beads were then washed 5 times with NT2 buffer, resuspended in 2x Laemmli buffer, and boiled at 95 degrees Celsius for 10 minutes. The supernatant was then stored at −80 degrees until they were used for western blots.

### *In vitro* whole-cell patch clamp recordings of human iPSC-derived neurons and mouse cortical neurons

Human iPSC-derived neurons were used for electrophysiology experiments between days 21-24 of the neural differentiation protocol. Whole-cell patch-clamp recordings were performed at room temperature using Multiclamp 700B amplifier (Molecular Devices) from borosilicate patch electrodes (P-97 puller and P-1000 puller; Sutter Instruments) containing a potassium-based intracellular solution (123 mM K-gluconate, 10 mM KCl, 10 mM HEPES; 1 mM EGTA, 1 mM MgCl2, 0.1 mM CaCl2, 1 mM Mg-ATP, and 0.2mM Na4GTP, pH 7.2). 0.06% sulpharhodamine dye was added to the intracellular solution to confirm the selection of multipolar neurons. The extracellular solution consisted of 140 mM NaCl, 5 mM KCl, 1.25 mM NaH_2_PO_4_, 1 mM MgCl_2_, 2 mM CaCl_2_, 10 mM glucose, and 10 mM HEPES (pH 7.4). Data was digitized at 10 – 20 kHz and low-pass filtered at 1 - 2 kHz. Recordings were omitted if access resistance was >30 MΩ. Whole-cell recordings were clamped at −70 mV and corrected for a calculated - 10mV junction potential. Rheobase was determined by a step protocol with 5 pA increments, where the injected current had 25 ms duration. Action potential waveform parameters were all analyzed in reference to the threshold. Repetitive firing step protocols ranged from -20 pA to +50 pA with 5 pA increments. No more than two neurons per coverslip were used to reduce the variability. Data were analyzed using the Clampfit software (Molecular Devices), while phase-plane plots were generated in the OriginPro software (Origin Lab). For GRIA1 overexpression experiments, mouse neurons were infected with GRIA1, GRIA1 R208H, and GRIA1 A636T lentiviral constructs at DIV11 and recorded on DIV14-15. Mouse neurons for electrophysiology experiments were cultured in Neurobasal media (supplemented with an additional 0.3 % (w/v) glucose and 0.22 % (w/v) NaCl). The same intracellular solution was used as the human neuron recordings, with a mouse artificial cerebrospinal fluid extracellular solution (125 mM NaCl, 2.5 mM KCl, 2 mM CaCl2, 1 mM MgCl_2_, 5 mM HEPES, 33 mM Glucose, pH 7.2). Whole-cell recordings of mouse neurons were clamped at −80 mV and corrected for a calculated −10mV junction potential.

### Staining and imaging of mouse cortical neurons and human iPSC-derived neurons

Mouse cortical neurons and human iPSC-derived neurons on coverslips were fixed with 4% paraformaldehyde (PFA) for 10 minutes at room temperature, washed once with PBS, and stored in PBS at 4 °C protected from light. Fixed coverslips were then blocked and permeabilized in BP solution (PBS with 10% donkey serum and 0.3% Triton-X) for 45 minutes at room temperature. Coverslips were then incubated with primary antibodies at 4 °C overnight. The following day, coverslips were washed three times with PBS for eight minutes each. Coverslips were then incubated with secondary antibodies for one hour at room temperature, followed by three washes with PBS. For human iPSC-derived neuron Synapsin1 staining, coverslips were incubated with 300 mM of DAPI for 15 minutes, before the third wash with PBS. Excess liquid was then removed from the coverslips and they were mounted onto VistaVision glass microscope slides (VWR) with 10 µL of Prolong Gold Anti-Fade mounting medium (Life Technologies).

For TOMM20 staining, mouse neurons were fixed with 4% PFA at 37°C for 10 min and then permeabilized with 0.5 % Triton X-100 for 10 minutes. Non-specific binding was blocked by incubation with 5 % donkey serum in PBS for 50 minutes at room temperature, followed by primary antibody incubation. The secondary antibody was added for 50 minutes at room temperature. Primary and secondary antibodies were diluted in PBS with 0.5 % BSA, 2.5 % Donkey-serum, and 0.15 % Triton X-100. After primary and secondary antibody incubation, three washing steps with PBS were performed. Then, coverslips were incubated with Phalloidin-488, for F-actin labeling, and Hoechst dye for 45 minutes at room temperature followed by three PBS washes. Coverslips were mounted onto slides using Fluoromount-G® (Southern Biotech) and were stored protected from light. Synapsin1 and BioID2 stained images were taken on the Zeiss LSM 700 confocal microscope with63x or 40x oil objective, respectively. Mito-dsRed images were taken on the Echo Revolve microscope with a 20x objective.

For Duolink Proximity Labeling assays, the manufacturers protocol was followed in detail (Sigma). DIV17 mouse neuron cultures were fixed in 4% PFA for 20 minutes, permeabilized and blocked as described above, and incubated with the primary antibody overnight at 4 degrees Celsius. Anti-Mouse MINUS and Anti-Rabbit PLUS probes were used with the FarRed In Situ detection reagents to identify co-localized proteins. Anti-MAP2 was co-stained on coverslips during the primary and secondary incubations. Coverslips were mounted with the DuoLink In Situe Mounting Medium with DAPI.

### Synapsin1 puncta analysis in human iPSC-derived neurons

Synapsin1 stained images were processed and analyzed with ImageJ software. The Synapsin1 antibody was co-immunostained with MAP2 to determine dendrites with presynaptic puncta. Five biological replicates, which represent five separate neural inductions, were used for synaptic analysis. 5-10 neurons per genotype per replicate were used. Imaging settings were kept the same between images and synapsin1 images were analyzed at the same threshold. Dendrites were traced using ImageJ and the measure tool was used to quantify the number and size of the puncta within the traced region.

### Proteomic profiling of *Taok2* KO mice cortical post-synaptic density fraction through LC-MS/MS

The right cortical lobes of three P21-23 *Taok2* KO mice and five P21-23 wildtype littermates were harvested and differential centrifugation was used to obtain the crude post-synaptic density fraction(Kwan et al., 2016). PSD fractionations were validated by western blot for PSD-95 and synaptophysin (data not shown). Final post-synaptic density pellets were resuspended using 8 M urea and 100 mM ammonium bicarbonate. Protein samples were then reduced with 10 mM Tris(2-carboxyethyl)phosphine for 45 min at 37 °C, alkylated with 20 mM iodoacetamide for 45 min at room temperature, and digested by trypsin (Promega) (1:50 enzyme-to-protein ratio) overnight at 37 °C. The peptides were desalted with the 10 mg SOLA C18 Plates (Thermo Scientific), dried, and labeled with Multiplex 10-plex TMT labels (Thermo) in 100 mM triethylammonium bicarbonate, and quenched with 5% hydroxylamine before combined. 40 μg of the pooled sample was separated into 60 fractions by high-pH reverse-phase liquid chromatography (RPLC) using a homemade C18 column (200 μm × 30 cm bed volume, Waters BEH 130 5 μm resin) running a 70 min gradient from 11 to 32% acetonitrile− 20 mM ammonium formate (pH 10) at a flow rate of 5 μL/min. Each fraction was then loaded onto a homemade trap column (200 μm × 5 cm bed volume) packed with POROS 10R2 10 μm resin (Applied Biosystems), followed by a homemade analytical column (50 μm × 50 cm bed volume) packed with Reprosil-Pur 120 C18-AQ 5 μm particles (Dr. Maisch) with an integrated Picofrit nanospray emitter (New Objective). LC-MS experiments were performed on a Thermo Fisher Ultimate 3000 RSLCNano UPLC system that ran a 3 h gradient (11− 38% acetonitrile−0.1% formic acid) at 70 nL/min coupled to a Thermo QExactive HF quadrupole-Orbitrap mass spectrometer. A parent ion scan was performed using a resolving power of 120 000; then, up to 30 of the most intense peaks were selected for MS/MS (minimum ion counts of 1000 for activation) using higher energy collision-induced dissociation (HCD) fragmentation. Dynamic exclusion was activated such that MS/MS of the same m/z (within a range of 10 ppm; exclusion list size = 500) detected twice within 5 seconds was excluded from the analysis for 30 seconds. Data were analyzed using Proteome Discoverer 2.2 (Thermo). For protein identification, search was against the SwissProt mouse proteome database (55,366 protein isoform entries), while the search parameters specified a parent ion mass tolerance of 10ppm, and an MS/MS fragment ion tolerance of 0.02Da, with up to two missed cleavages allowed for trypsin. Dynamic modification of +16@M was allowed. Only proteins with two unique peptides were used for further analysis. Differentially expressed proteins (DEPs) were calculated through Significance B outlier test using the Perseus software(Tyanova et al., 2016), and only proteins that had adj. p-value < 0.05 were considered as DEPs.

### Transcriptome profiling of *Taok2* KO mice cortices through RNA sequencing

Cortices from *Taok2* KO and wildtype littermates were also harvested for RNA at post-natal day 21-23, with 3 males and 3 females from each genotype. The RNA was extracted using Trizol and was sent for total RNA sequencing at the Center for Applied Genomics (TCAG). mRNA was purified using poly(A) selection to avoid contamination of ribosomal RNAs and miRNAs. All samples were run on one lane resulting in ∼31-34 million of read pairs per sample. All analysis was carried out using the open-source platform Galaxy (usegalaxy.org)(Afgan et al., 2018). RNA reads were checked for good quality using the FastQC tool. The trimmomatic tool was used to identify and trim off known adaptors and remove any bases that have a Phred score of less than 20. FastQC was used again to ensure that adaptor sequences were removed and that the quality of the reads was not affected. We next used the HISAT2 alignment program for alignment of the RNA sequences to the mouse genome GRCm38 (NCBI). On average 85% of reads from mouse samples were aligned once and 5% were aligned more than once to distinct genome locations. Moving on, the featureCounts tool was used to count the number of reads per gene using the same reference genome as the HISAT2 tool. The DESeq2 tool was used to determine the significant differentially expressed genes (DEGs) between *Taok2* WT and KO mouse cortices. Genes were considered as DEGs if they had an adjusted p-value lower than 0.05.

### Gene set enrichment analysis (GSEA) of *Taok2* KO mouse proteome and transcriptome

DEGs and DEPs were ranked based on the equation –log_10_(adj. p-value)*Ln(fold change). GSEA 4.1.0 (Broad Institute)(Daly et al., 2003; Subramanian et al., 2005) was used to run the GSEA preranked test. Tests were run with 1000 permutations, weighted enrichment statistics, and excluding gene sets smaller than 15 and larger than 500 genes. All other settings were kept as default. All mouse GO term gene sets without electronic GO annotations (http://download.baderlab.org/EM_Genesets/) were used for the analysis(Merico et al., 2010). Visualization of the enriched gene sets was done on Cytoscape 3.8.2 using the EnrichmentMap app and the AutoAnnotate app was used for clustering similar gene sets(Bader et al., 2014; Kucera et al., 2016; Shannon et al., 2003). All visualized gene sets had an FDR < 0.1.

### Seahorse assay of *in vitro* mouse and human iPSC-derived neurons

Mouse cortical neurons were cultured as described above at a density of 30,000 cells/well in the Seahorse XF96 cell culture microplate. CRISPR/Cas9 KO mouse neurons were infected at DIV 7 and assayed at DIV 14. Human iPSC-derived NGN2 neurons were plated on day 4 of dox induction at a density of 50,000 cells per well in the Seahorse XF96 cell culture microplates (Agilent), pre-coated with 20 µg/ml Laminin and 67 µg/ml Poly-ornithine (Sigma). Mouse glia was plated on top of the neurons at a density of 25,000 cells per well, 24 hours later. Cells were used for the Seahorse assay on day 7. The day prior to the seahorse assay the Seahorse XFe96 sensor cartridge was filled with Calibrant XF solution and incubated at 37 °C (without CO_2_) overnight. On the day of the assay the Seahorse XF96 microplates were washed twice with 200 µl per well of pre-warmed MST media (Seahorse XF DMEM pH 7.4 media supplemented with 1 mM sodium pyruvate, 2.5 mM GlutaMAX, and 17.5 mM Glucose). The plate was then filled with 180 µl per well MST media and incubated at 37 °C (without CO2) for 1 hour. During the incubation, the mitochondrial stress test drugs were added to the XFe96 sensor cartridge (1 µM Oligomycin for mouse neurons and 3 µM Oligomycin for human neurons, 1 µM FCCP, and 1 µM Rotenone/Antimycin A resuspended in MST media). The cartridge plate with the drug compounds were then put in the Seahorse XFe96 analyzer for calibration. After calibration, the microplate was placed into the Seahorse XFe96 analyzer for the pre-set mitochondrial stress test protocol. Oxygen consumption rates (OCR) were recorded every seven minutes and the drug compounds were added in 21-minute intervals. Oligomycin was used to inhibit ATP-synthase to measure ATP-synthase dependant respiration, FCCP was added to decouple the inner membrane to measure maximal respiration, and Rotenone and Antimycin A were added together to measure non-mitochondrial respiration. After the assay, microplates were frozen at − 80°C overnight and cell content was measured using the Cyquant cell proliferation assay (Thermo) by measuring fluorescence with the CLARIOStar machine and MARS data analysis software (BMG LABTECH). Cellular respiration analysis was performed using the Wave software (Agilent) and OCR values were normalized to the number of cells per well.

### CRISPR/Cas9 knockout in mouse cortical neurons

Mouse cortical neurons were infected at DIV7 with the pLV CMV-Cas9-T2A-EGFP (MOI 1) and pLV U6-sgRNA/EF1a-mCherry (MOI 3) lentiviruses. Cultures were allowed to recover until DIV14 and were then taken for the seahorse assay. The GeneArt genomic cleavage detection kit (Thermo) was used to detect insertions or deletions in the targeted sites.

### Measuring mitochondrial activity in shRNA knockdown mouse neurons

Embryonic age E15 C57BL6/J mouse pups were *in utero* electroporated with Taok2 shRNA and control shRNA. Electroporated mouse embryo cortices were than harvested and cultured at E18. Mouse neuron cultures were imaged at DIV5 after incubation with 2nM TMRM (Thermo). Images were analyzed on ImageJ. Soma regions were delineated and integrated density in the soma (soma area x mean intensity) was measured. For background correction, mean background intensity was obtained from the neighbouring region.

### Measuring mitochondrial activity and content in mouse cortical neurons

DIV 6 mouse cortical neurons cultured from *Taok2* WT, Het, and KO mouse embryos were incubated with 2 nM TMRM (Invitrogen, #T668) and/or 100 nM MitoTracker Green (Cell Signaling Technology, #9074P) were directly added to the conditioned medium, and incubated for 15 minutes. Cells were then imaged within 30 minutes after the incubation time. Images were loaded onto ImageJ, background mean intensity was measured from the region without TMRM and MitoTracker signals inside the cell, then the cell was delineated and the background was removed. After background correction, using the JACoP plugin the TMRM-MitoTracker signal colocalization was analyzed using Manders’ correlation coefficients. For Manders’ correlation coefficients, threshold values for TMRM (red channel) and MitoTracker (green channel) were set to 335±55 and 640±50 respectively. 16-bit wide field images were taken on a Nikon EclipseTi2 inverted spinning disk microscope equipped with 60X oil (NA 1.4) objective, an LED light source (Lumencor® from AHF analysentechnik AG, Germany), and a digital CMOS camera (ORCA-Flash4.0 V3 C13440-20CU from Hamamatsu) controlled with NIS-Elements software. The microscope imaging chamber is equipped to maintain 37 °C temperature and 5 % CO_2_. Illumination, exposure and gain settings were kept the same across different conditions for imaging TMRM and MitoTracker signals.

### TOMM20 staining analysis for mitochondria content in mouse cortical neurons

DIV 7-8 mouse cortical neurons cultured from *Taok2* WT, Het, and KO mouse embryos were fixed and stained for TOMM20. 16-bit Z-series images with a step size of 300 nm Images were acquired on confocal spinning disk microscope using a 60X oil (NA 1.4) objectives. Illumination, exposure and gain settings were kept the same across the conditions. The images were loaded onto ImageJ and z-projection (sum slices) for the entire cell in z-axis was performed on the confocal images. Using ImageJ, soma region was carefully delineated and total intensity, also known as integrated density, in the soma (soma area * mean intensity) was measured. For background correction, mean intensity (background mean intensity) was obtained from the neighbouring region (out of the cell). Using the following equation, we obtained the corrected values. Corrected value = total intensity in the soma – (background mean intensity * soma area).

### Electron microscopy of synaptic mitochondria from mouse brain cortices

Coronal vibratome sections of the cingulate cortex (cg1 and cg2) and the prelimbic cortex (PL) of the PFC, the primary somatosensory regions S1HL, S1Fl, S1BF, and the intermediate HC were collected and prepared for electron microscopy as described in Richter *et al*.(Richter et al., 2019). Semithin sections (0.5 µm) were prepared for light microscopy mounted on glass slides and stained for 1 min with 1% Toluidine blue. Ultrathin sections (60 nm) were examined in an EM902 (Zeiss, Munich, Germany). Pictures were taken with a MegaViewIII digital camera (A. Tröndle, Moorenweis, Germany). EM images that were collected and analyzed for synapse formation on the dendritic spines or shafts from Richter *et al*. were reanalyzed for mitochondrial morphology. Mitochondria morphology from the EM images obtained from Taok2 Wt and Taok2 KO genotypes were analyzed manually using ImageJ. based on their morphology the mitochondria are and categorized to Category 1 - Normal mitochondria with well stacked Cristae, Category 2 - mitochondria with enlarged Cristae, Category 3 - mitochondria with condensed Cristae.

### Mito7-dsRed puncta analysis in human iPSC-derived neurons

*TAOK2* KO, A135P and wildtype human iPSC-derived NGN2 neurons were transfected with 0.8 µg of Mito7-dsRed (Addgene #55838) and 0.2 µg of pCAG-Venus at day 5, with 2 µl of Lipofectamine 2000 (Thermo). Venus was used to trace neuron projections.10 neurons per genotype from two separate neural inductions were used for analysis. Imaging settings were kept the same between images and Mito7-dsRed images were analyzed at the same threshold. Dendrites were traced using ImageJ and the measure tool was used to quantify the size of the puncta within the traced region

### Correlation of 41 ASD-risk gene PPI networks

Corrplot (R package) was used to create the correlation plot. The normalized biotinylation score to the bait protein was used to calculate the correlation between ASD-risk gene PPI networks. The Silhouette and Within cluster sum of squares methods were used to calculate the optimal kmeans number for clustering. Genes were ordered by hierarchal clustering.

### Cell type/DEG/ASD gene list enrichment analysis

Human cell type gene expression and ASD DEGs and ASD gene lists were obtained from their respective publications(Feliciano et al., 2019; Nowakowski et al., 2017; Ruzzo et al., 2019; Sanders et al., 2015; Satterstrom et al., 2020; Velmeshev et al., 2019; Wilfert et al., 2021; Yuen et al., 2017). For the enrichment analysis we used the Fisher exact test comparing each gene list with the shared ASD-risk gene PPI network in the mouse brain background protein list, which was used for pathway enrichment analysis. P-values and ODDs ratios were calculated for each comparison. To account for multiple comparisons, Bonferroni correction thresholds were calculated as p = 0.05 divided by the number of comparisons.

### Clinical score analysis

Rare variants of individuals diagnosed with ASD were extracted from the MSSNG database (research.mss.ng)(Yuen et al., 2017). Only variants with estimated high or medium impact strengths were used for analysis, and variants were categorized into three groups (missense variants, splicing variants, and frame shift/premature stop codon variants). Adaptive behavior and socialization standard scores of affected individuals were extracted from the MSSNG associated Metabase (data-explorer.mss.ng). Individuals were grouped based on the presence of mutations in the 41 ASD-risk genes that were clustered into three groups. Individuals that had variants in genes between multiple groups were not included in the analysis. Separate analyses were carried out between individuals grouped by missense, splicing or frame shift/premature stop codon variants. Clinical data was considered as non-parametric and the Kruskal-Wallis ranked test with post hoc Dunn’s test was used for comparison between the adaptive behavior and socialization standard scores of each group.

### Data representation and figure generation

Networks and gene set enrichment maps were created on Cytoscape v3.8.2. Graphs were created on GraphPad Prism 7. Representative electrophysiology traces were extracted onto CorelDRAW. Microscopy images were prepared using ImageJ. Dot plots, correlation plots, and heat maps were created on R Studio. Graphical abstract and flowcharts were created on and exported from BioRender.com (BU24JATEEX, SD235B8ORF, KW235KT7TM, RZ235KTA0S). Final figures were organized and created using Adobe Illustrator CC.

## QUANTIFICATION AND STATISTICAL ANALYSIS

Data are expressed as mean ± s.e.m, except the clinical analysis which is shown as a box and whisker plot showing the minimum, median, and maximum scores. A minimum of three biological replicates were used for all experiments, where separate HEK cell transfections, iPSC dox-inductions, mouse neuron cultures, or littermates are considered as individual biological replicates. All statistical analysis was done on GraphPad Prism 7. All comparisons were assumed to be parametric, except for the clinical score analyses. ROUT’s outlier test was used to identify possible outliers, with a Q value of 0.1 %. For statistical analysis unpaired t-test, or One-Way ANOVA and Two-Way ANOVA with *post hoc* Holm-Sidak tests were used to compare all experimental conditions to the control condition. All unpaired t-tests were two-sided, except for the one-sided t-test used for identification of BioID2 prey proteins. Clinical scores were assumed to be non-parametric and the Kruskal-Wallis H test with post hoc Dunn’s test was used to compare all groups to each other. Any variation from the described statistical analyses is described and explained in the figure legends. The p-values are defined in the figure legends and p < 0.05 are considered statistically significant.

## Supplementary Tables

**Table S1.** BioID2 PPI networks of 41 ASD-risk genes and cellular compartment genes. Related to Figure 2.

**Table S2.** Comparison of BioID2 PPI networks identified in HEK293 cells and mouse cortical neurons. Related to Figures 1, S2 and S4.

**Table S3.** Comparison of BioID2 PPI network enriched cellular components identified in HEK293 cells and mouse cortical neurons. Related to Figures 1, S3 and S4.

**Table S4.** BioID2 PPI network enriched cellular components of compartment specific genes. Related to Figure 2.

**Table S5.** BioID2 PPI network enriched pathways of 41 ASD-risk genes. Related to Figure 2.

**Table S6.** 41 ASD-risk gene PPI network map enriched pathways. Related to Figure 2.

**Table S7.** Differentially expressed genes and proteins and dysregulated pathways in *Taok2* KO mouse cortices. Related to Figures 3 and S7.

**Table S8.** Comparison of BioID2 PPI networks between ASD-risk genes and their variants. Related to Figure 4.

**Table S9.** BioID2 PPI network enriched pathways of ASD-risk genes and their variants. Related to Figure 4.

**Table S10.** List of sources for 41 ASD-risk genes and cellular compartment genes. Related to Figure 1.

