## Supplementary Figures for "Neuron-specific protein network mapping of autism risk genes identifies shared biological mechanisms and disease relevant pathologies"

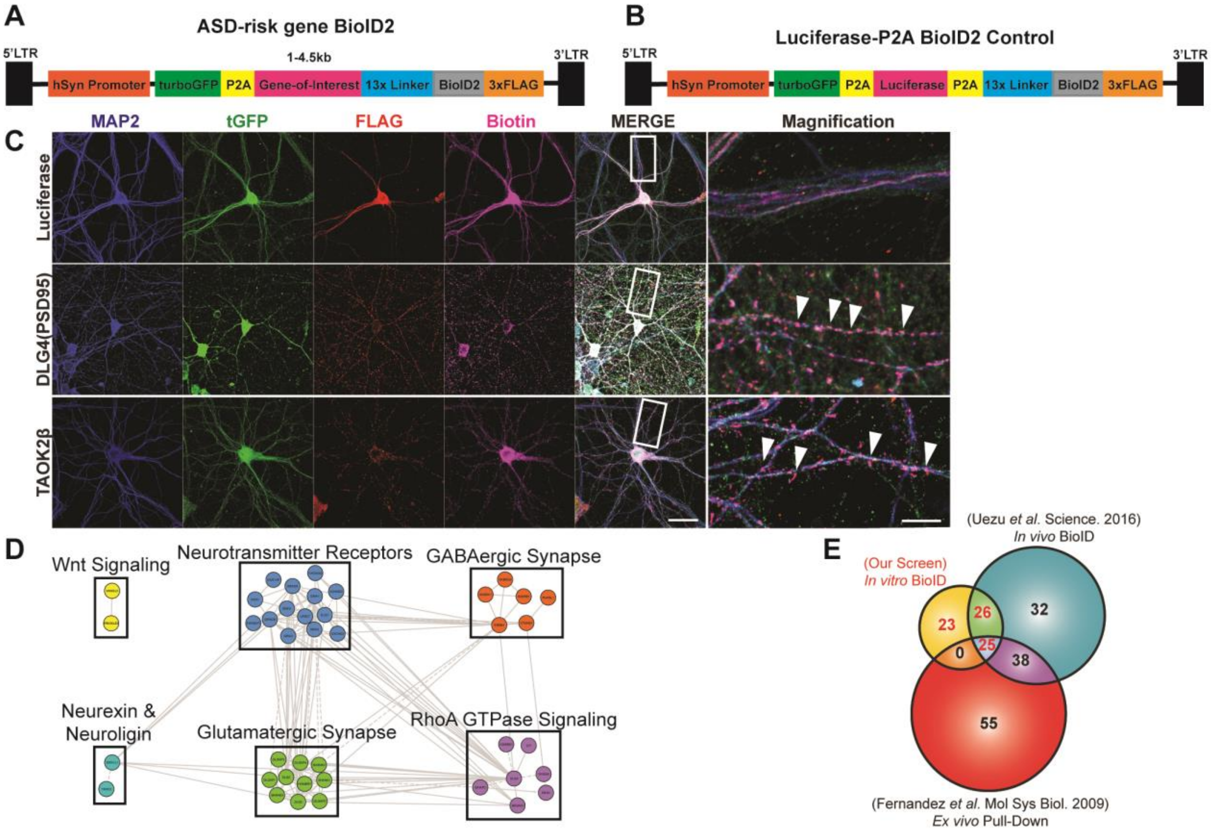


**Figure S1. BioID2 of DLG4 in mouse cortical neurons.** (A) Diagram of the BioID2 fusion construct for the 41 ASD-risk genes. (B) Diagram of the control Luciferase control fusion construct. (C) Representative images of cortical neurons infected with the Luciferase-P2A-BioID2, PSD95-BioID2, and TAOK2β-BioID2 constructs. Scale bar is 20μm. Magnified images are shown on the right. White arrows point to synaptic localization of PSD95-BioID2. Scale bar is 5μm. (D) Reactome pathways enriched in the PSD95 PPI network. Clusters created using the Reactome FI app on Cytoscape and labeled with most significantly enriched pathways for each cluster (adj. $p<0.05$). (E) Venn diagram of shared protein interactors between our in vitro PSD95 PPI network and proteins identified by mouse in vivo PSD95 BioID (Uezu et al. 2016) or mouse in vivo tandem affinity purification of PSD95 (Fernandez et al. 2009). Related to Figure 1.

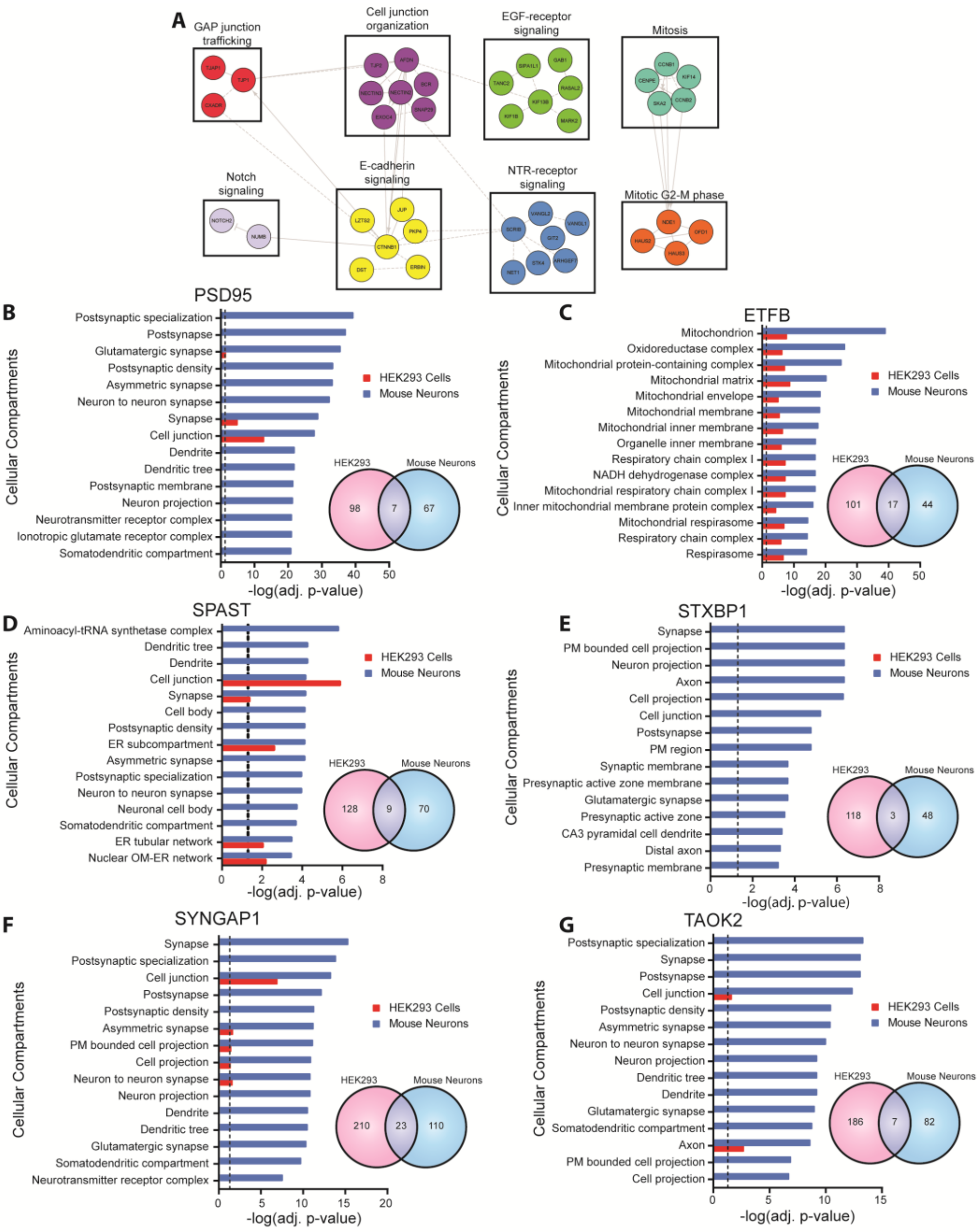


**Figure S2. Comparison of ASD-risk gene PPI networks from BioID2 in HEK293 cells and mouse cortical neurons.** (A) Reactome pathways enriched in the PSD95 PPI network from HEK293 cells. Clusters created using the Reactome FI app on Cytoscape and labeled with the most significantly enriched pathways (adj. p<0.05). Top 15 cellular compartments from mouse cortical neurons (blue) enriched in the PPI networks of PSD95 (B), ETFB (C), SPAST (D), STXBP1 (E), SYNGAP1 (F), and TAOK2 (G) compared to enrichment in HEK293 cells (red) (g:Profiler, Benjamini-Hochberg FDR adj. p<0.05). Adjacent Venn diagrams show shared protein interactors identified by BioID2 in HEK293 cells vs mouse cortical neurons. PM: Plasma membrane, OR: Outer membrane, ER: endoplasmic reticulum. Related to Figure 1.

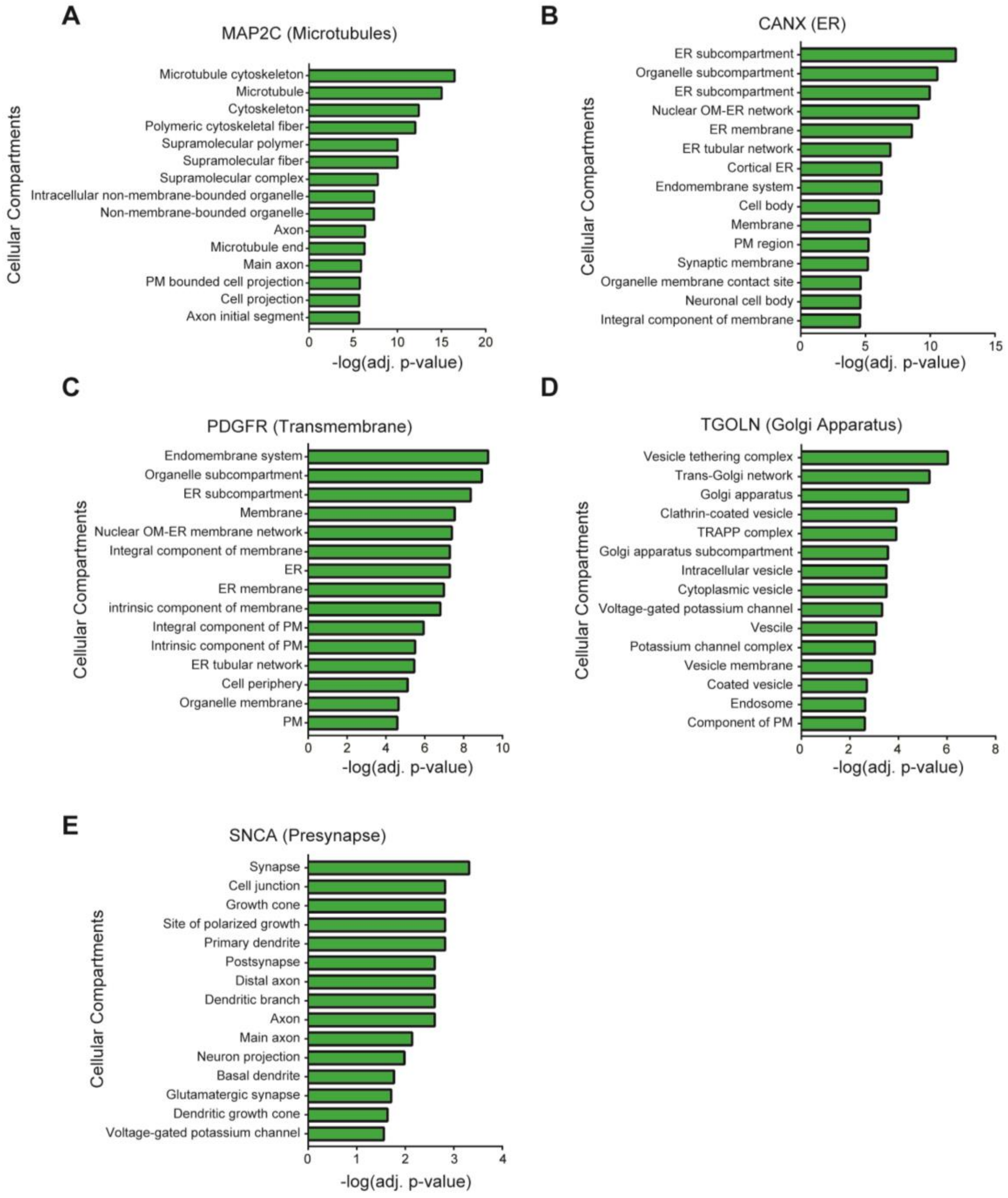


**Figure S3. Validation of BioID2 using genes localized to specific compartments in mouse cortical neurons.** BioID2 of cellular compartment proteins MAP2C (A), CANX (B), PDGFR (transmembrane domain) (C), TGOLN (D), and SNCA (E). g:Profiler pathway enrichment was used to identify significantly enriched cellular compartments (g:Profiler, Benjamini-Hochberg FDR adj. $p<0.05$). PM: plasma membrane, OM: outer membrane, ER: endoplasmic reticulum. Related to Figure 1.

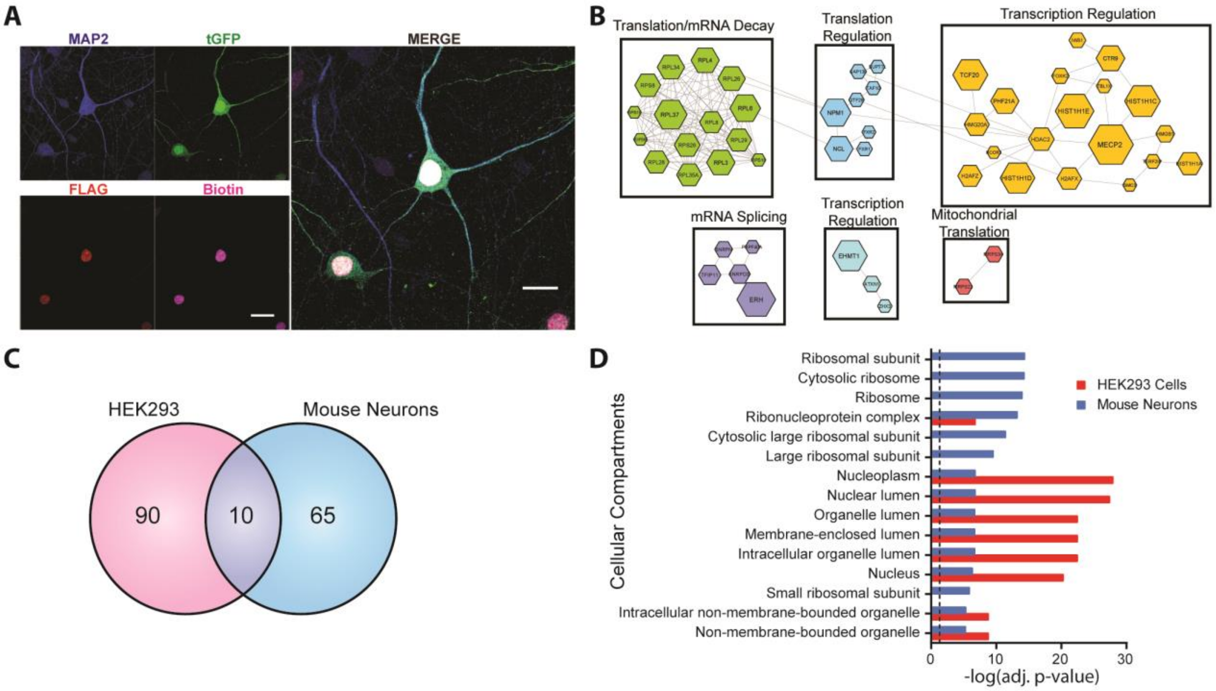


**Figure S4. BioID2 of MECP2 in mouse cortical neurons.** (A) Representative images of cortical neurons infected with the MECP2-BioID2 construct. Scale bar is 20μm. (B) Reactome pathways enriched in the MECP2 PPI network. Clusters created using the Reactome FI app on Cytoscape and labeled with most significantly enriched pathways (adj. p<0.05. (C) Venn diagram shows shared protein interactors identified by BioID2 in HEK293 cells vs mouse cortical neurons. (D) Top 15 cellular compartments from mouse cortical neurons (blue) enriched in the MECP2 PPI network compared to enrichment in HEK293 cells (red) (g:Profiler, Benjamini-Hochberg FDR adj. p<0.05). Related to Figure 1.

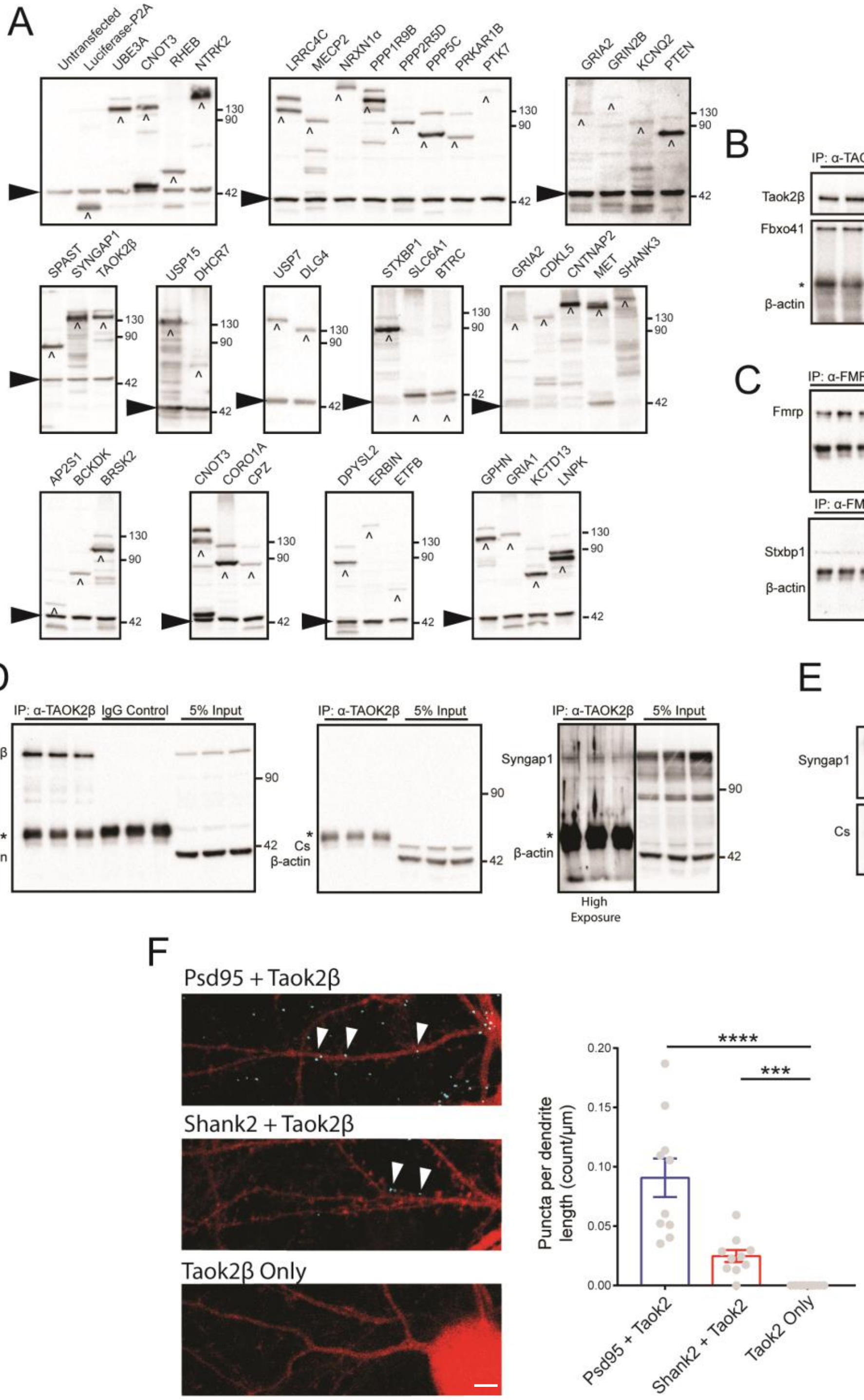


**Figure S5. Western blots of 41 ASD-risk gene BioID2 constructs.** (A) Western blots of ASD-risk gene BioID2 constructs transfected in HEK293 cells and immunoblotted for FLAG and β-actin as the loading control. ^ denotes expected BioID2 fusion protein size. Bands higher than the caret indicated bands are possible tGFP fusion proteins due to P2A inefficiency. Arrow denotes β-actin loading control. Bands lower than β-actin are possible degraded BioID2-FLAG proteins. (B) Coimmunoprecipitation of Fbxo41 with Taok2β from three separate CD1 mouse cortices. (C) Coimmunoprecipitation of Stxbp1 with Fmrp from three separate CD1 mouse cortices. (D) Coimmunoprecipitation of Syngap1 with Taok2β from three separate CD1 mouse cortices. Center blot was stripped and reblotted for Syngap1 (*right*). (E) Coimmunoprecipitation of Syngap1 with Citrate synthase from three separate CD1 mouse cortices. * denotes presence of heavy and light chain proteins. (F) Representative images of DIV17 mouse cortical neurons infected with mCherry showing co-localization of Psd95 and Taok2β, and Shank2 and Taok2β (*right*) using PLA. Scale bar is 5μm. Quantification showing increased number of puncta in mouse cortical neurons compared to Taok2β alone (*left*). 10 randomly selected dendrites from 10 neurons per condition; two separate mouse neuron cultures. Mean ± s.e.m. ***$p<0.001$, ****$p<0.0001$. Related to Figure 2.

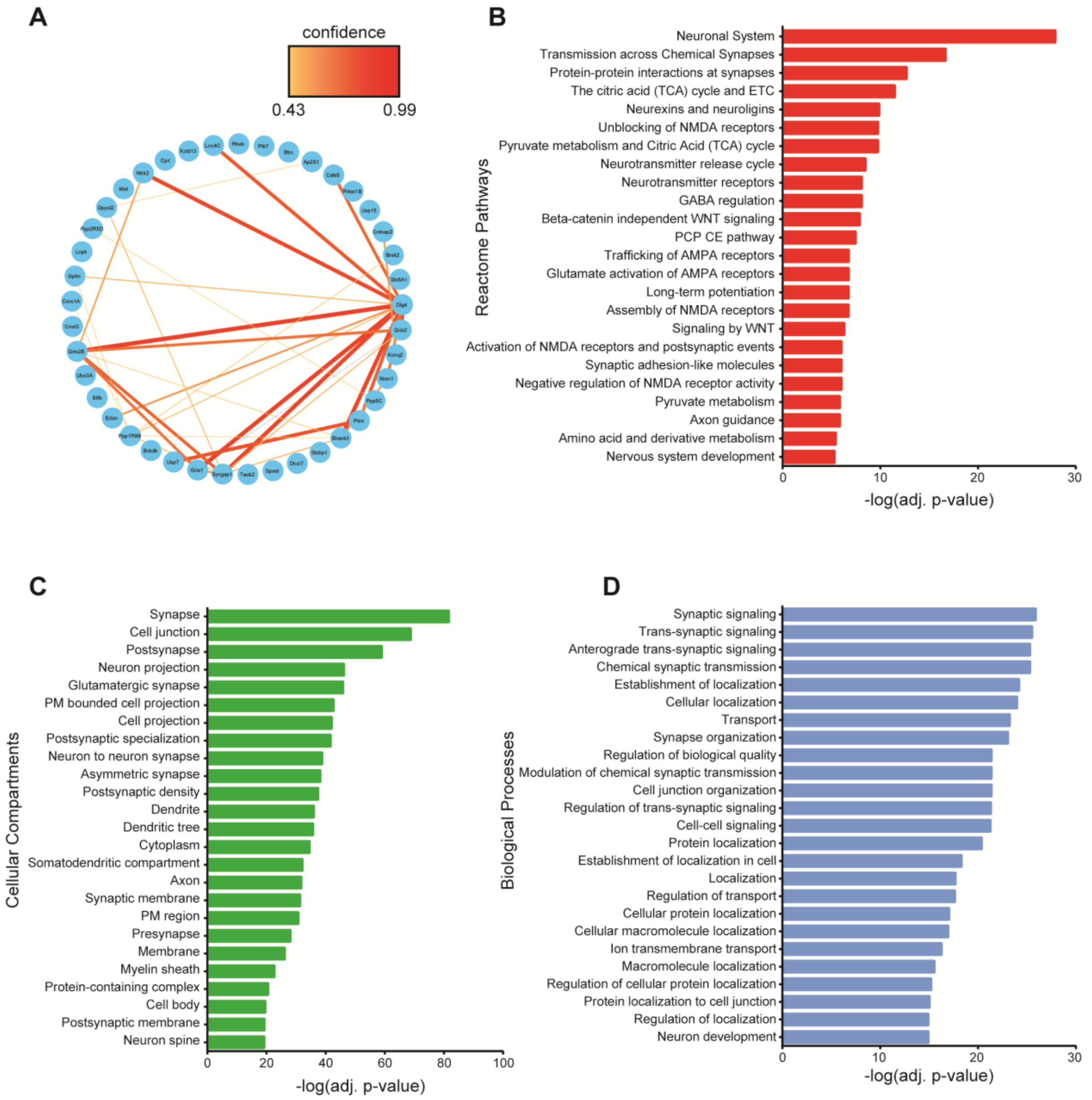


**Figure S6. Enriched pathways in the shared ASD-risk gene PPI network map identified using BioID2.** (A) Known physical interactions of the 41 ASD-risk genes from the STRING database. Color and increasing thickness of the line represents the confidence of the interaction starting at medium interaction confidence (0.4). Pathway enrichment was used to identify significantly enriched Reactome pathways (B), cellular compartments (C), and biological processes (D) (g:Profiler, Benjamini-Hochberg FDR adj. p<0.05). The top 25 pathways are shown for each graph. PM: plasma membrane, OM: outer membrane, ER: endoplasmic reticulum. Related to Figure 2.

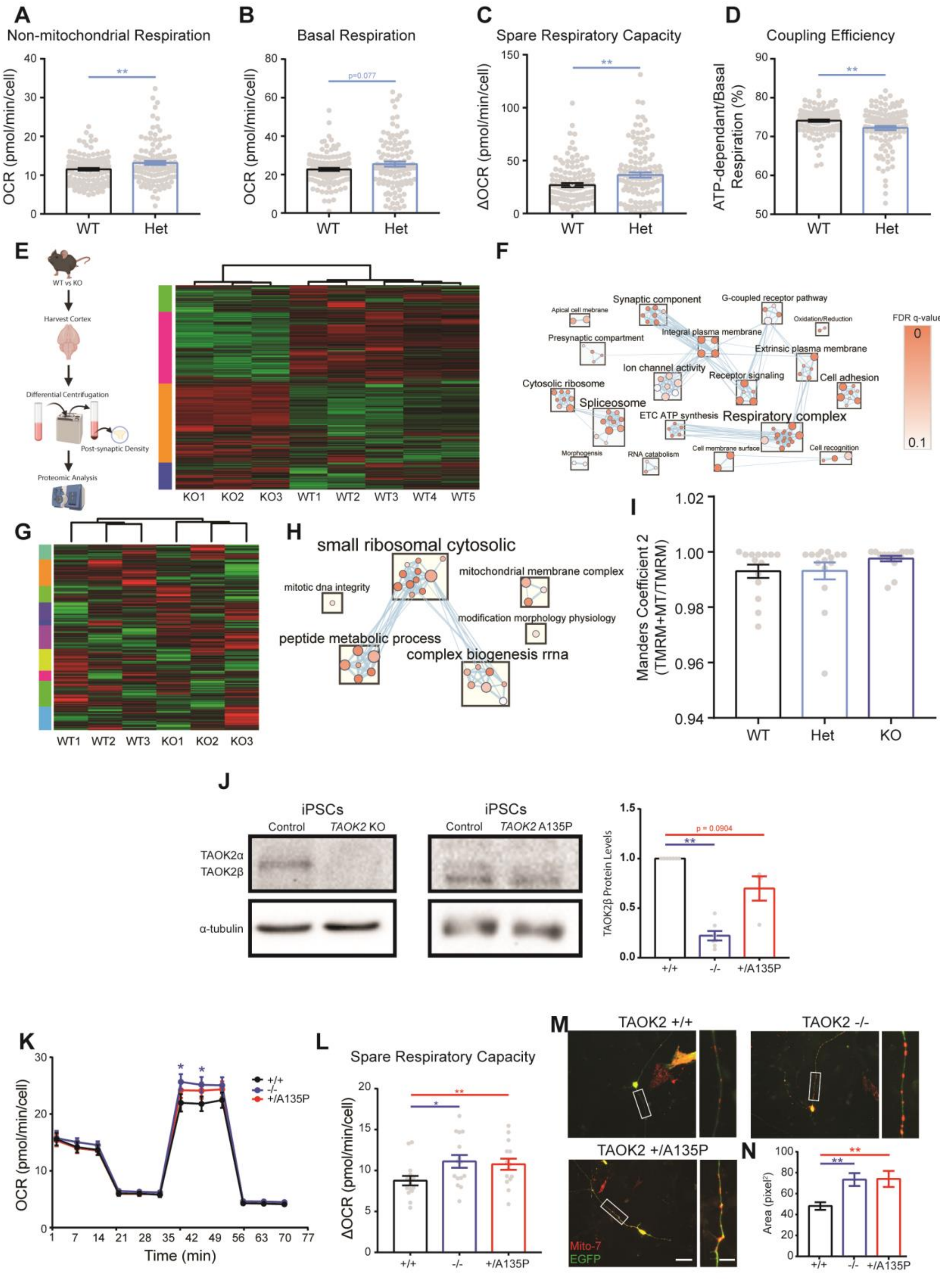
A
Non-mitochondrial Respiration
OCR (pmol/min/cell)
WT
Het
B
Basal Respiration
p=0.077
C
Spare Respiratory Capacity
ΔOCR (pmol/min/cell)
D
Coupling Efficiency
ATP-dependant/Basal Respiration (%)
E
WT vs KO
Harvest Cortex
Differential Centrifugation
Post-synaptic Density
Proteomic Analysis
KO1 KO2 KO3 WT1 WT2 WT3 WT4 WT5
F
Apical cell membrane
Synaptic component
G-coupled receptor pathway
Oxidation/Reduction
Presynaptic compartment
Integral plasma membrane
Extrinsic plasma membrane
Ion channel activity
Receptor signaling
Cell adhesion
Cytosolic ribosome
Spliceosome
ETC ATP synthesis
Respiratory complex
Morphogensis
RNA catabolism
Cell membrane surface
Cell recognition
FDR q-value
G
WT1 WT2 WT3 KO1 KO2 KO3
H
small ribosomal cytosolic
mitotic dna integrity
mitochondrial membrane complex
modification morphology physiology
peptide metabolic process
complex biogenesis rrna
I
Manders Coefficient 2 (TMRM+MT/TMRM)
WT
Het
KO
J
iPSCs
Control
TAOK2 KO
TAOK2 A135P
TAOK2α
TAOK2β
α-tubulin
TAOK2β Protein Levels
p = 0.0904
+/+
-/-
+/A135P
K
OCR (pmol/min/cell)
Time (min)
L
Spare Respiratory Capacity
ΔOCR (pmol/min/cell)
M
TAOK2 +/+
TAOK2 -/-
TAOK2 +/A135P
Mito-7
EGFP
N
Area (pixel²)

**Figure S7. Loss or disruption of ASD-risk gene TAOK2 causes alterations in cellular respiration and mitochondrial proteins.** (A-D) Taok2 Het neurons have increased non-mitochondrial respiration and spare respiratory capacity, and decreased coupling efficiency. WT = 118 wells, Het = 122 wells from three separate cultures. (E) Shotgun proteomics of post-synaptic density fraction from Taok2 WT and KO mouse cortices. (F) Taok2 KO PSD fractions have significant decrease in synaptic and mitochondrial protein gene sets (GSEA, FDR<0.1.; five Taok2 WT and three Taok2 KO mice littermates). Size of nodes represents number of proteins and color represents FDR q-value. (G) RNA sequencing of Taok2 WT and KO mouse cortices. (H) TAOK2 KO mouse cortices have altered mRNA levels of mitochondrial membrane proteins. (GSEA, FDR<0.1, three Taok2 WT and KO mice littermates each). Size of nodes represents number of proteins and color represents FDR q-value. (I), All active mitochondria are stained by MitoTracker. WT = 14, Het = 15, KO = 15 neurons from three separate cultures). (J) Western blot of CRISPR/Cas9-edited iPSCs and neurons showing loss of TAOK2 expression in TAOK2 KO (-/-) and A135P (+/A135P) lines. WT = 7, KO = 7, A135P = 4 wells from separate iPSC cultures. (K-L) DIV7 TAOK2 KO human neurons have significantly increased maximal respiration. WT = 118 wells and Het = 122 wells from three separate cultures). TAOK2 KO and A135P neurons have significantly increased spare respiratory capacity. WT = 118 wells, Het = 122 wells from three separate cultures). (M) Representative images of human neurons transfected with Mito7-DsRed constructs at DIV7 and fixed and imaged at DIV9. Scale bar is 20 µm. Magnification of boxed areas shown on the right. Scale bar is 5 µm. (N) TAOK2 KO and A135P neurons have larger Mito7-DsRed punctae size compared to wildtype neurons. WT = 71, KO = 520,  A135P = 421 punctae from 15-16 neurons per genotype). Mean ± s.e.m. *p<0.05, **p<0.01. Related to Figure 3.

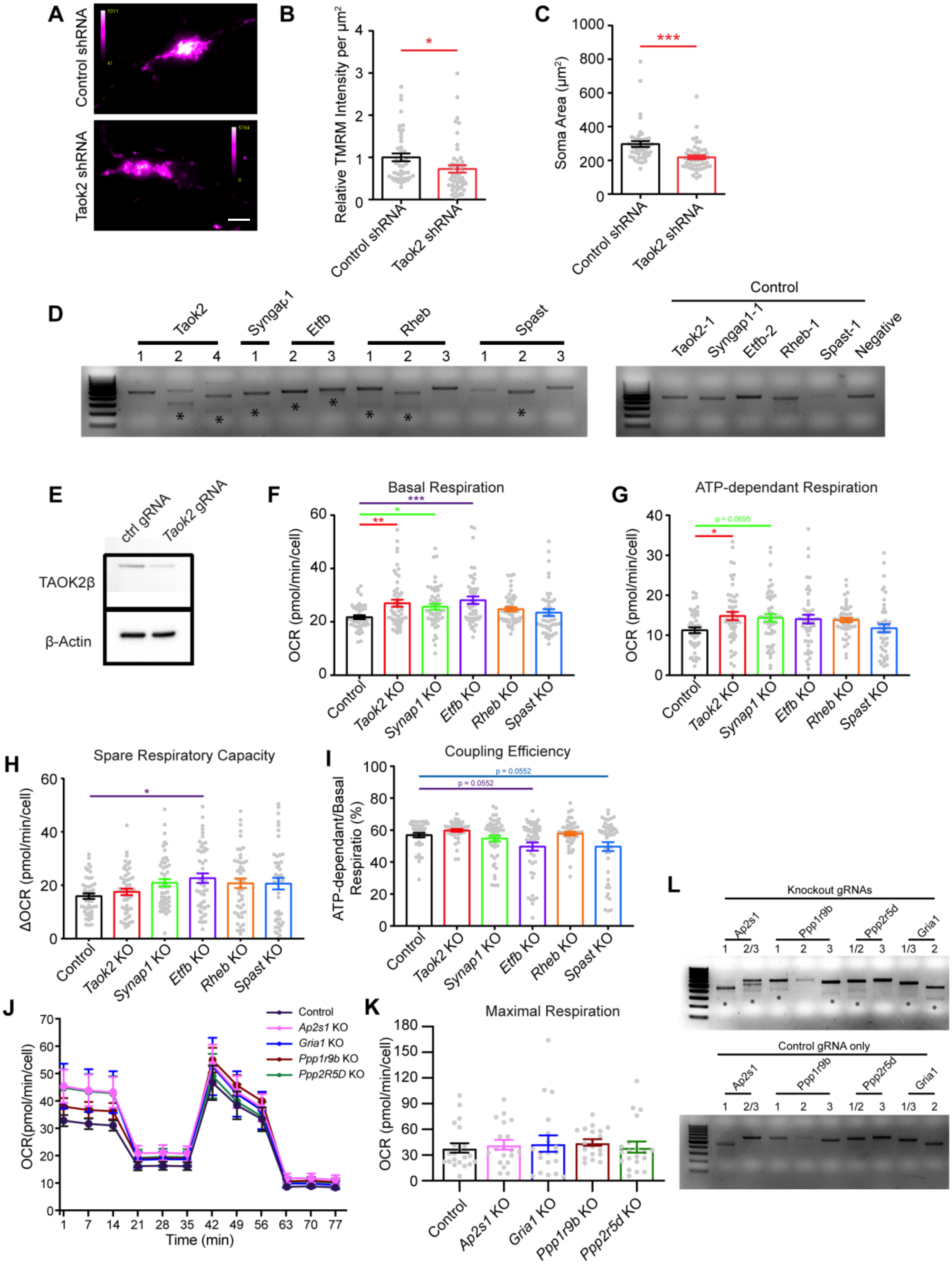
A
Control shRNA
Taok2 shRNA
B
Relative TMRM Intensity per µm²
Control shRNA
Taok2 shRNA
C
Soma Area (µm²)
Control shRNA
Taok2 shRNA
D
Taok2
Syngap1
Etfb
Rheb
Spast
Control
Taok2-1
Syngap1-1
Etfb-2
Rheb-1
Spast-1
Negative
E
ctrl gRNA
Taok2 gRNA
TAOK2β
β-Actin
F
Basal Respiration
OCR (pmol/min/cell)
G
ATP-dependant Respiration
p = 0.0695
OCR (pmol/min/cell)
H
Spare Respiratory Capacity
ΔOCR (pmol/min/cell)
I
Coupling Efficiency
p = 0.0552
p = 0.0552
ATP-dependant/Basal Respiratio (%)
Control
Taok2 KO
Synap1 KO
Etfb KO
Rheb KO
Spast KO
J
Control
Ap2s1 KO
Gria1 KO
Ppp1r9b KO
Ppp2R5D KO
OCR(pmol/min/cell)
Time (min)
K
Maximal Respiration
OCR (pmol/min/cell)
Control
Ap2s1 KO
Gria1 KO
Ppp1r9b KO
Ppp2r5d KO
L
Knockout gRNAs
Control gRNA only
Ap2s1
Ppp1r9b
Ppp2r5d
Gria1

**Figure S8. Acute knockout of ASD-risk genes in mouse cortical neurons alters mitochondrial activity and cellular respiration.** (A) Representative images of mouse cortical neurons infected with control shRNA or Taok2 shRNA showing TMRM intensity. Scale bar is 10μm. (B) Mouse cortical neurons with acute knockout of Taok2 have decreased relative TMRM activity, even with decreased soma size (C). WT = 44, Taok2 KD = 49 neurons from two separate cultures. (D) Indel cleavage assay for Taok2, Syngap1, Etfb, Rheb, and Spast KO shows at least at least one disrupted gRNA target region in mouse cortical neurons. *Indicates secondary band due to digested indel. (E) Reduced Taok2 protein expression in mouse cortical neurons infected with Cas9 and Taok2 KO gRNAs. Infected at DIV7 and harvested at DIV18 for western blot of TAOK2β. Significant changes in different aspects of cellular respiration (Basal respiration (F),ATP- dependent respiration (G), spare respiratory capacity (H), and coupling efficiency (I) in mouse cortical neurons with CRISPR/Cas9 KO of Taok2, Syngap1,or Etfb. Taok2 KO = 51 wells, Syngap1 KO = 50 wells, Etfb KO = 47 wells, Rheb KO = 48 wells, Spast KO = 45 wells from five separate cultures). (J) CRISPR/Cas9 KO of *Ap2s1*, *Gria1*, *Ppp1r9b*, and *Ppp2r5d* show no changes in cellular respiration or maximal respiration (K) in DIV14 mouse cortical neurons. Control = 21 wells, Ap2s1 KO = 19 wells, Gria1 KO = 19 wells, Ppp1r9b KO = 19 wells, Ppp2r5d KO = 20 wells from five separate cultures). (L) Indel cleavage assay for Ap2s1, Gria1, Ppp1r9b, Ppp2r5d KO shows at least at least one disrupted gRNA target region in mouse cortical neurons. *Indicates secondary band due to digested indel. Mean ± s.e.m. *$p<0.05$, **$p<0.01$, ***$p<0.001$.  Related to Figure 3.

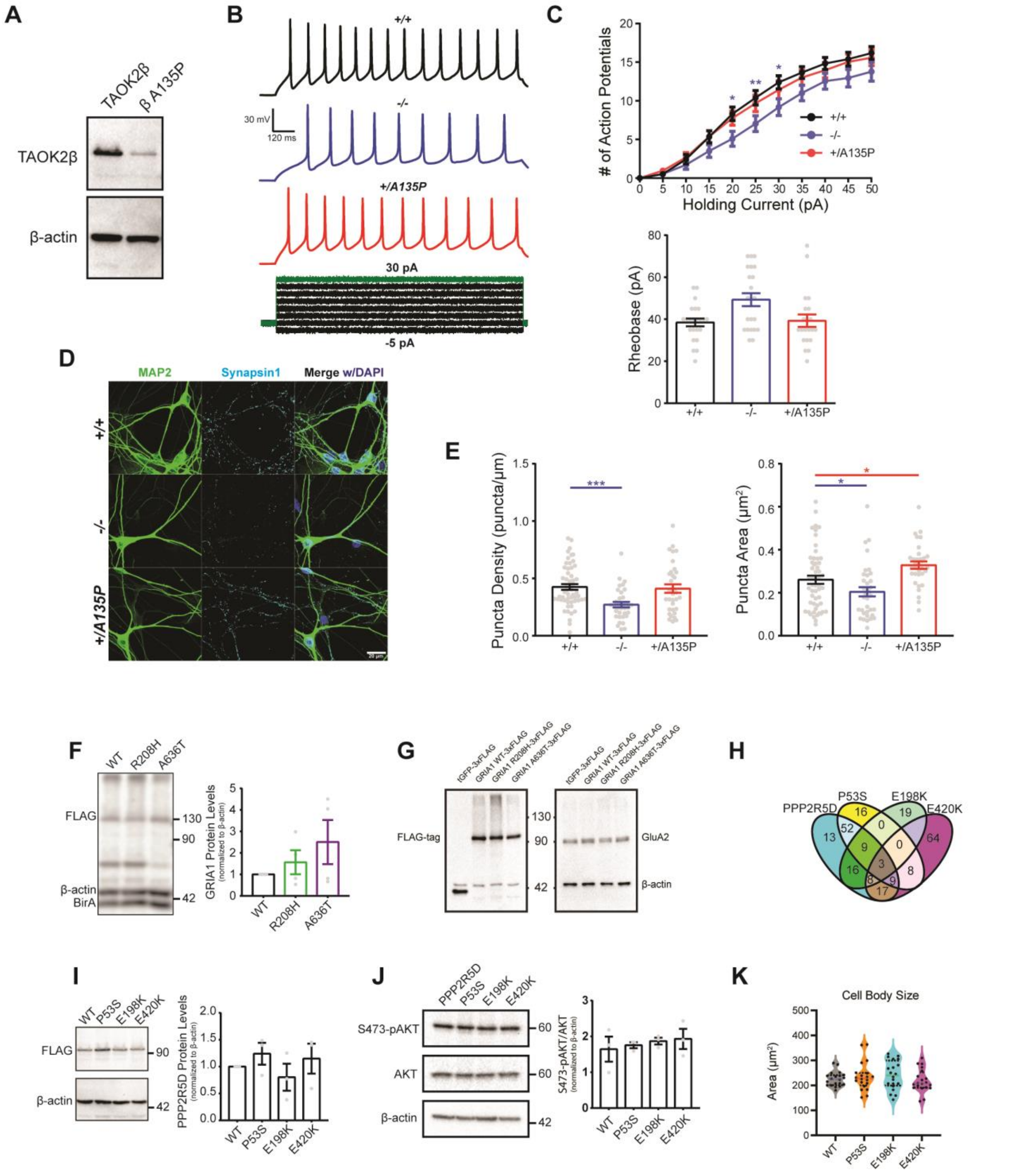


**Figure S9. De novo mutations in TAOK2 caused altered synaptic transmission and neuron firing.** (A) Western blot of TAOK2 WT and A135P BioID2 constructs in HEK293 cells. (B) Representative traces of repetitive firing (*top*) and current injection (*bottom*). (C) TAOK2 KO neurons have reduced repetitive firing (*top*) and increased rheobase (*bottom*). WT = 23, KO = 22, A135P = 21 neurons from 3 separate transductions. (D) Representative images of TAOK2 WT (+/+), KO, and A135P human neurons, stained with MAP2, Synapsin 1, and DAPI 21 days after NGN2 induction. (E) Reduced synapsin punctae density and size in TAOK2 KO neurons (*left*) and increased synapsin puncta size in TAOK2 A135P neurons (*right*). WT = 55, KO = 35, A135P = 34 neurons from five separate transductions. (F) Representative western blot of GRIA1 WT, R208H, and A636T BioID2 constructs expressed in HEK293 cells (*left*) and quantification (*right*) showing no significant difference. Four separate transfections. (G) Western blot of GluA2 subunit in DIV16 cortical neurons infected with GRIA1 WT, R208H, and A636T-FLAG lentiviral constructs for 4 days (H) Venn diagram of PPI network proteins of PPP2R5D WT and variants. (I) Representative western blot of PPP2R5D WT, P53S, E198K, and E420K BioID2 constructs expressed in HEK293 cells (*left*) and quantification (*right*) showing no difference in expression. Three separate transfections. (J) Representative western blot of p-AKT and AKT in HEK293 cells expressing PPP2R5D WT or variant constructs expressed in HEK293 cells (*left*) and quantification (*right*) showing no difference in the pAK/AKT ratio. Three separate transfections. Quantification of western blots in (E), (H), and (I) were done with the volume intensity of the Flag-tag band. (K) Neurons expressing the PPP2R5D variants show no changes in neuron size. 20 neurons from 5 separate infections per condition. Mean ± s.e.m. *$p<0.05$, **$p<0.01$, ***$p<0.001$. Related to Figure 4.

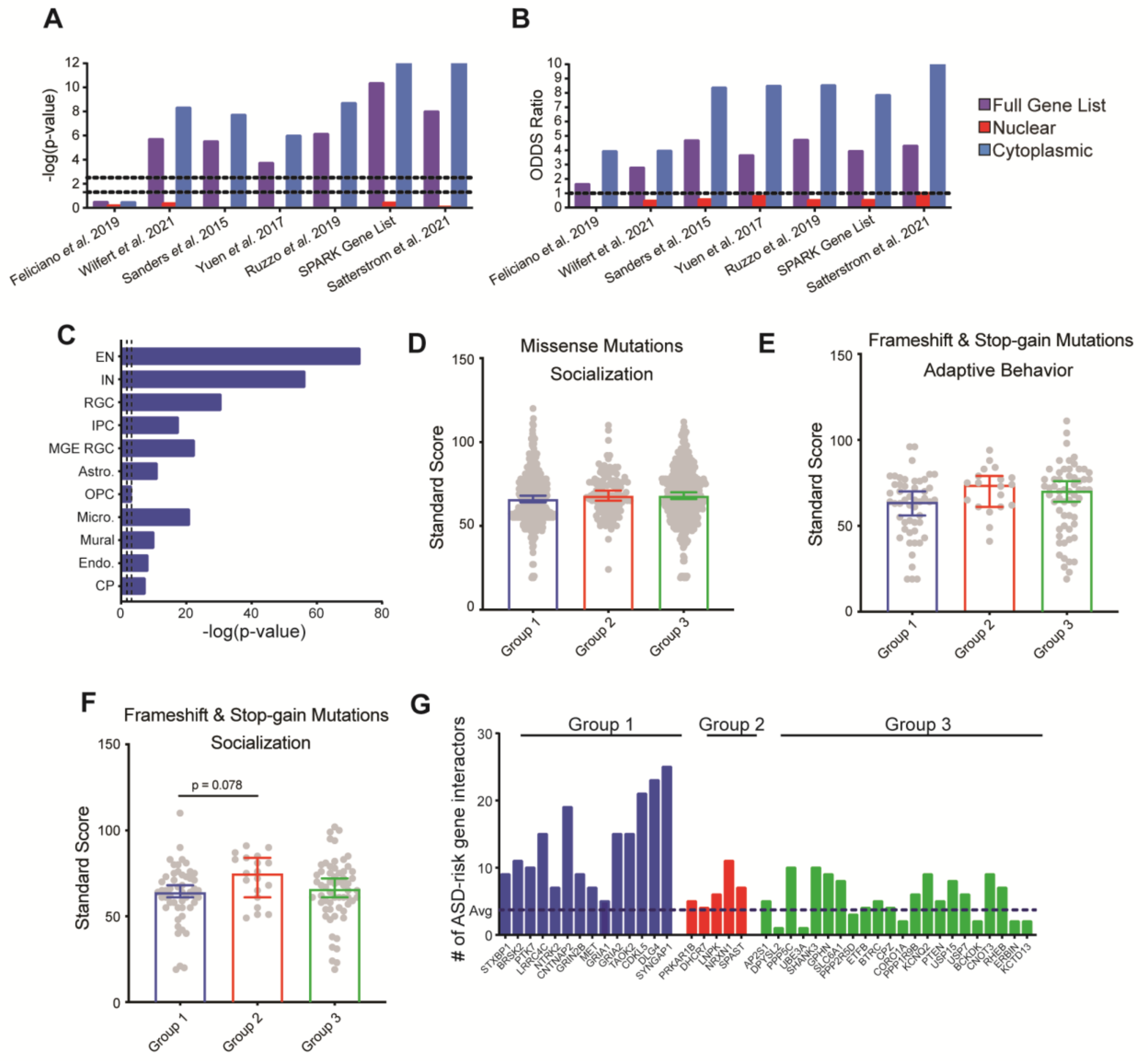


**Figure S10. Enrichment of ASD risk-genes in the shared ASD PPI network map and grouping of clinical phenotypes.** (A) Enrichment of full gene list, cytoplasmic gene only lists, and nuclear gene only lists from published works and SPARK, in the shared ASD-risk gene PPI network shown by signficance. (Fisher's exact test). Dashed lines represent nominal (p = 0.05, *left*) and Bonferroni corrected (p = 0.05/number of cell types, *right*) significance thresholds. (B) ODDs ratio of full gene list, cytoplasmic gene only lists, and nuclear gene only lists enriched in the shared ASD-risk gene PPI network. (C) The shared ASD-risk gene PPI network enriches for human neuron cell types (Fisher's exact test). Dashed lines represent nominal (p = 0.05, *left*) and Bonferroni corrected (p = 0.05/number of cell types, *right*) significance thresholds. EN = excitatory neurons, IN = inhibitory neurons, RGC = radial glial cells, MGE RGC = medial ganglionic eminence, IPC = intermediate progenitor cells, Astro. = astrocyte, OPC = oligodendrocyte progenitor cells, Micro. = microglia, Endo. = endothelial cells, CP = choroid plexus cells, C-F = cortico-fugal, C-C = cortico-cortico, PV = paravalbumin, SST = somatostatin, VIP = vasoactive intestinal peptide, FB = fibrous, PP = protoplasmic, Neu_NRGN = neurogranin-expressing. Light blue bars have nominal p-value significance, while dark blue bars have Bonferroni corrected significance. (D) Individuals with missense mutations in Cluster 1, 2 and 3 genes show no significant differences in socialization standard scores (Non-parametric Kruskal-Wallis test, p = 0.1765, post-hoc Dunn's test; Group 1 = 351, Group 2 = 114, and Group 3 = 416 probands). Individuals with frameshift or stop gain mutations in Cluster 1, 2 and 3 genes show no significant differences in adaptive behavior (E) and socialization standard (F) scores (Non-parametric Kruskal-Wallis test, adaptive behaviour: p = 0.1069, Group 1 =51, Group 2 = 19, and Group 3 = 60 probands; socialization: p = 0.0803, Group 1 = 51, Group 2 = 19, and Group 3 = 60 probands; post hoc Dunn's test). (G) Number of ASD-risk genes identified in each of the 41 ASD-risk gene protein-protein interactions. Dashed line represents average expected risk genes. Box and whisker plot (minimum, 1st quartile, median, 3rd quartile, maximum). *p<0.05, **p<0.01. Related to Figure 5.
